# Tissue-wide scRNA-seq analysis reveals enrichment of imprinted genes in stem and endocrine cell-types in mice

**DOI:** 10.1101/2023.08.05.552036

**Authors:** Matthew J. Higgs, Matthew J. Hill, Rosalind M. John, Anthony R. Isles

## Abstract

Enriched expression of imprinted genes may provide evidence of convergent function. Here we interrogated five single-cell RNA sequencing datasets to identify imprinted gene over-representation in the embryonic and adult mouse focusing on tissues including the bladder, pancreas, mammary gland and muscle. We identify a consistent enrichment of imprinted genes in stromal cell and mesenchymal stem cell populations across these tissues, suggesting a role in tissue maintenance. Furthermore, we identify a distinct enrichment in the endocrine islets of the mouse pancreas, over and above the stromal/stem cells from this tissue. Taken together with our previous work examining imprinted gene expression in cell subpopulations of the adult mouse brain and pituitary gland, these data suggest that genomic imprinting influences physiology largely via separate systems of cell populations either involved in hormonal signalling or in stemness and cell-fate co-ordination.

## INTRODUCTION

Imprinted genes are expressed preferentially (often exclusively) from one allele according to their parent of origin via epigenetic regulation (John et al., 2022). These genes have been shown to be key regulators of embryonic and neonatal growth and development (Ferguson-Smith, 2011) and, accordingly, are implicated in a number of childhood growth disorders (Butler, 2009, Angulo et al., 2015, Eggermann et al., 2021, Eggermann et al., 2015). Since the discovery and identification of specific imprinted genes, much work has been done to characterise the function of individual genes. Current consensus supports the idea that imprinted genes regulate fetal growth, development and metabolism (Cassidy and Charalambous, 2018, Millership et al., 2019, Smith et al., 2006, Lambertini et al., 2012). However, with the identification of imprinted gene networks (Al Adhami et al., 2015, Gabory et al., 2009, Varrault et al., 2006), it has become increasingly valuable to think about and investigate imprinted genes as a gene set. This approach is a logical progression from theories on how imprinting evolved, which predict that imprinted genes will converge to affect specific developmental and metabolic processes important in mammals (Haig, 2000, Wolf and Hager, 2006).

This convergent role for imprinted genes - modulating fetal and postnatal growth, metabolism and resource acquisition - involves several embryonic and adult tissues for which selected imprinted genes have individually been implicated functionally. These tissues include the brain (Davies et al., 2008, Ho-Shing and Dulac, 2019, Wilkinson et al., 2007), the pancreas (Stefan et al., 2011, Millership et al., 2018, de Mora et al., 2003), the mammary gland (Hanin and Ferguson-Smith, 2020, Xu et al., 2020, Yonekura et al., 2019), adipose tissue (Takahashi et al., 2005, Curley et al., 2005), muscle tissue (Gabory et al., 2009, Yan et al., 2003) and extra-embryonic tissues/placenta (Coan et al., 2005, John, 2017, Lefebvre, 2012). This combination of endocrine and metabolic tissues co-ordinates appetite, metabolic rate, fat/lean mass and the behaviour of animals as offspring to acquire maternal resources (food, warmth etc. see John et al. (2023)). However, the ideas relating to which physiologies genomic imprinting impacts have often relied on findings from a selection of high-profile imprinted genes, and yet most recent estimates suggest the number of imprinted genes in the mouse genome is now over 260 (Tucci et al., 2019) and so whole genome approaches are also necessary to support the idea that imprinted genes converge on specific processes which might indicate their evolutionary purpose.

Previously, genome-wide approaches for imprinted genes have investigated allelic expression in order to identify tissues with the largest number of genes showing parent of origin biases. Several imprinted gene atlases have been produced by measuring allele specific expression from multiple tissues (Andergassen et al., 2017, Babak et al., 2015), and these tend to show the same pattern concerning tissue imprinting patterns. First, imprinted genes when expressed almost always show an allelic bias (Babak et al., 2015). Second, the brain, placenta and embryonic tissues possess the largest number of imprinted genes. Nevertheless, other tissues, such as the adrenal gland and pancreatic islets (Babak et al., 2015) and the mammary gland (Andergassen et al., 2017), express large numbers of imprinted genes. High numbers of imprinted gene expressed in a given tissue may indicate a physiological function of particular significance for imprinting, and many of these tissues align with the expected functions of imprinted genes. However, the total number of genes expressed within different tissues varies (Han et al., 2018). This means the number of imprinted genes expressed in a given tissue may reflect the nature of gene expression in that tissue, rather than reflecting a unique hub of imprinted activity. For instance, the brain expresses most of the genome (Lein et al., 2007, Negi and Guda, 2017) so it is perhaps not surprising that a large number of imprinted genes are expressed here. To negate this problem, we have applied an unbiased, statistical approach to identify brain regions and specific cell types within the brain enriched for imprinted genes (Higgs et al., 2022). We found convergent enrichment of imprinted genes in behaviourally- and endocrine-associated hypothalamic neurons and the mid- and hind-brain monoaminergic system. Furthermore, we have shown that the enriched genes identified in these analyses can have functional consequences when disrupted, using the example of parenting behaviour and the associated neurons (Higgs et al., 2023).

In this study we adopt the same method for identifying over-represented upregulation of imprinted genes and apply it to murine tissues at a single-cell resolution. Not only did this shed light on the cell types driving the enrichment of tissues found in our earlier multiple-organ analysis (Higgs et al., 2022) but it was informative to identify cell populations enriched in body tissues impacted by imprinted genes. We began by analysing general cell population enrichments from embryonic, neonatal and adult tissues before analysing several adult tissues in depth (Figure 1). This included previously identified tissues - pancreas, bladder and muscle tissues (Higgs et al., 2022) and the mammary gland based on over-represented cell types we saw in these analyses. We identify a consistent enrichment of imprinted genes in stromal cells and mesenchymal stem cell populations across various tissues, and a distinct signal from the endocrine islets of the mouse pancreas, over and above the stromal/stem cells from these tissues. These findings suggest a role for imprinted genes in tissue maintenance and, when taken together with our previous work examining imprinted gene expression in neuroendocrine cell subpopulations of the adult mouse brain and pituitary, the endocrine regulation of adult physiology.

**Figure 1.**
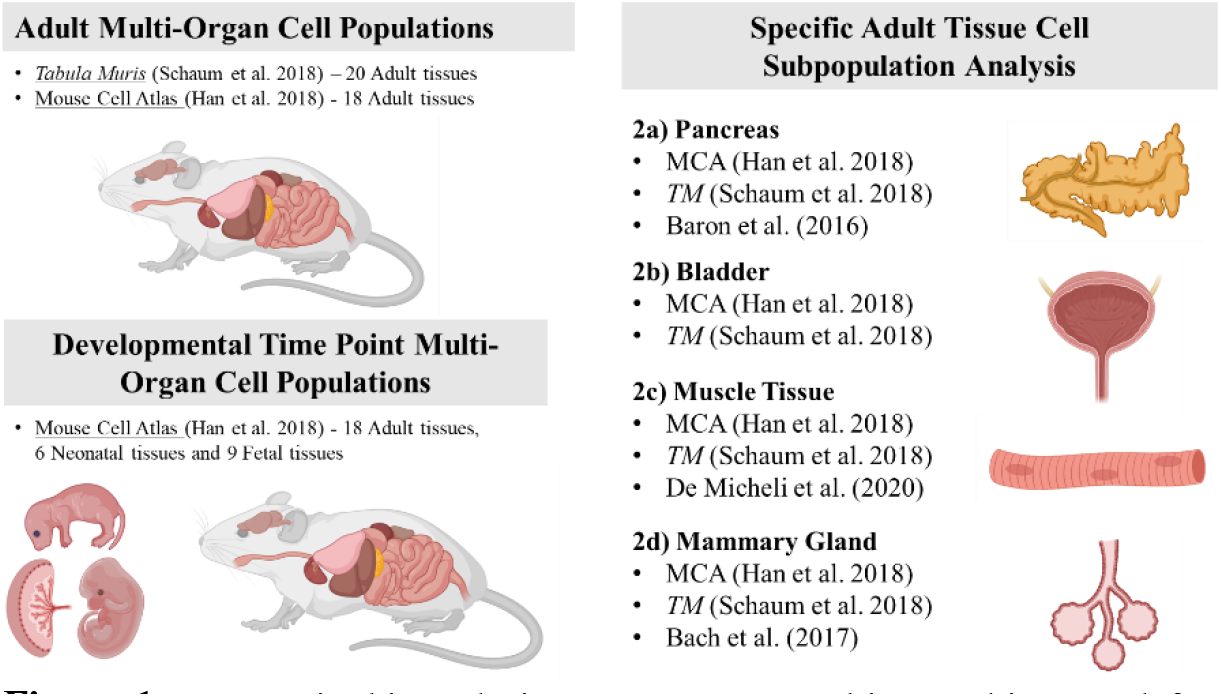
Datasets in this analysis. Datasets are sorted into multi-organ (left) and tissue-specific (right) analyses. The original publications are provided for each analysis.

## RESULTS

### Imprinted genes are over-represented in stromal and mesenchymal cell populations across multiple tissues

The Mouse Cell Atlas (MCA) (Han et al., 2018) and the Tabula Muris (TM) (Schaum et al., 2018) are single cell compendiums containing ∼20 overlapping, but not identical, adult mouse organs. Our previous analysis found imprinted gene over-representation in cells of the brain, pancreas, bladder and muscle tissues. However, we did not isolate cell populations within these tissues which could be driving this tissue enrichment. In the present study we carried out an over-representation analysis (ORA) on both datasets, grouping the cells by their cell subpopulation identities as classified by the original authors (e.g., ductal cell, excitatory neuron). All data were processed according to the original published procedure, a list of upregulated genes was produced for each cell/identity group (vs. all other cell/identity groups) and a one-sided Fisher’s Exact test was performed using a custom list of imprinted genes (Supplemental Table S1) to identify cell populations in which imprinted genes were over-represented amongst the upregulated genes for that cell population. Each dataset in this study was analysed independently which allowed us to look for convergent patterns of enrichment between datasets of similar tissues/cell-types.

Cells from the 18 adult tissues of the MCA were distinguished into 292 tissue-specific cell types. Cell-types with an over-representation of imprinted genes from the analysis of the adult MCA cells are shown in Table 1 (See Supplemental Table S2 for full output). Stromal cells from various organs (incl. pancreas, bladder and mammary gland) were found to be the major over-represented cell type. MCA stromal cells were identified by Han et al. (2018) as ‘connective tissue cells that support the function of parenchymal cells’ and were identified based on expression of collagens, laminins, elastin and fibronectin. This includes cells such as fibroblasts, extracellular matrix (ECM) and mesenchymal stem/stromal cells (Meirelles et al., 2006, Schäfer et al., 2012, Valkenburg et al., 2018). To identify situations in which imprinted genes were enriched amongst the stronger markers of a tissue/cell-type, we performed a Gene-Set Enrichment Analysis (GSEA) on tissues meeting minimum criteria (see Methods), which assessed whether imprinted genes were enriched within the top ranked upregulated genes for that tissue (ranked by Log2 Fold Change). No cell-types in this analysis showed a significant GSEA for imprinted genes.

**Table 1.**
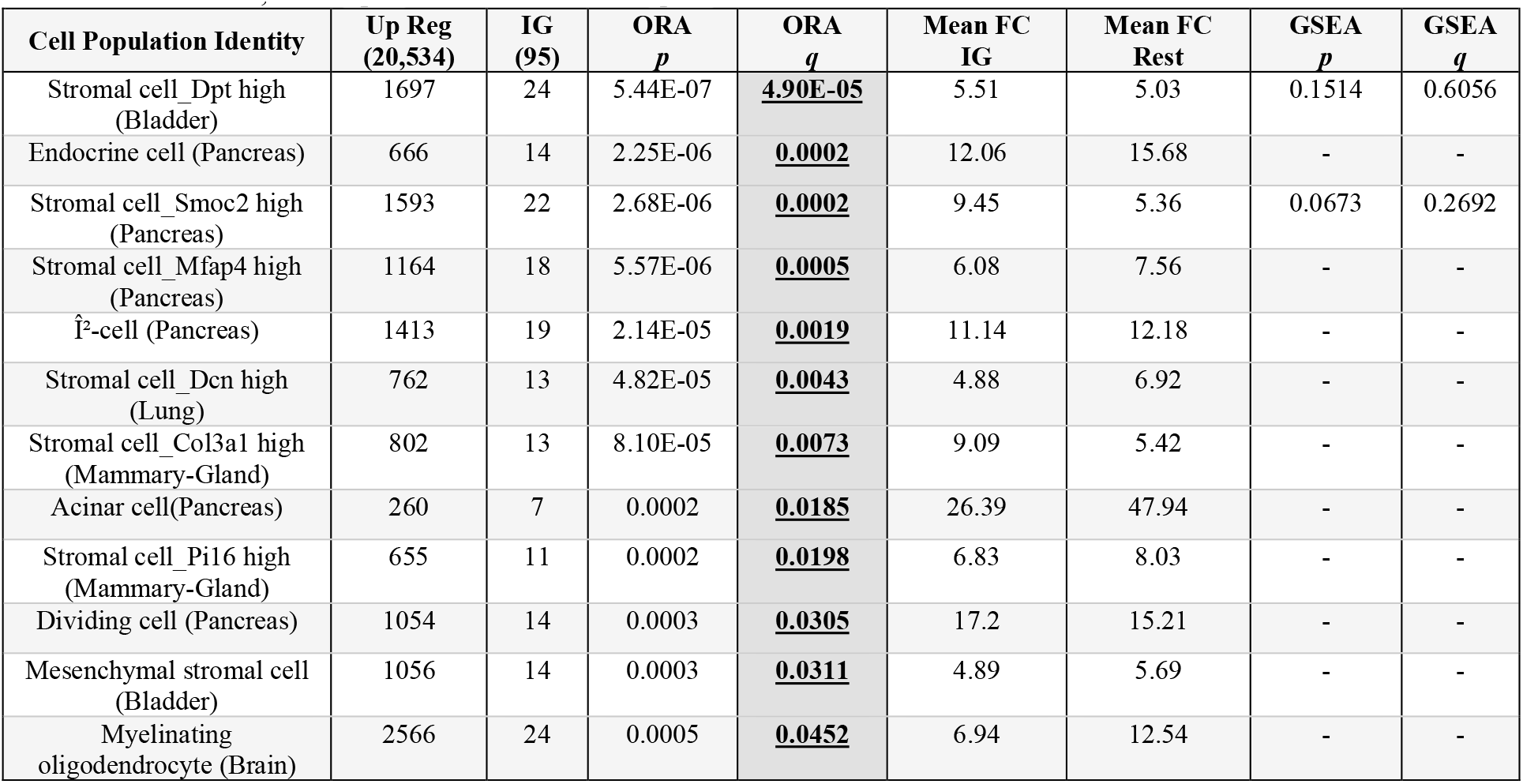
Imprinted gene over-representation in MCA adult cell populations (Han et al., 2018). Identity – Cell identities for the cells used in analysis; Up Reg – number of upregulated genes (total number of genes in the dataset in brackets number of imprinted genes upregulated with 2FC (total number of IGs in the dataset in brackets); ORA p – p value from over representation analysis on groups with minimum 5% of total IGs; ORA q – Bonferroni corrected p value from ORA; Mean FC IG – mean fold change for upregulated imprinted genes; Mean FC Rest – mean fold change for all other upregulated genes; GSEA p – p value from Gene Set Enrichment Analysis for identity groups with 15+ IGs and Mean FC IG > Mean FC Rest; GSEA q – Bonferroni corrected p values from GSEA.

Cells from the 20 tissues of the TM were distilled into 89 tissue-spanning cell types. Significant findings from the analysis of these cell types are shown in Table 2 (See Supplemental Table S3 for full output). Over-representation was predominantly seen in pancreatic and muscle-based cell types but cell types from the brain, bladder, mammary gland and adipose were also found over-represented from this large-scale comparison. Importantly, stromal cells, originating from the mammary gland and lung, and various mesenchymal stem cells were sites of enrichment, consistent with the results from the MCA.

**Table 2.**
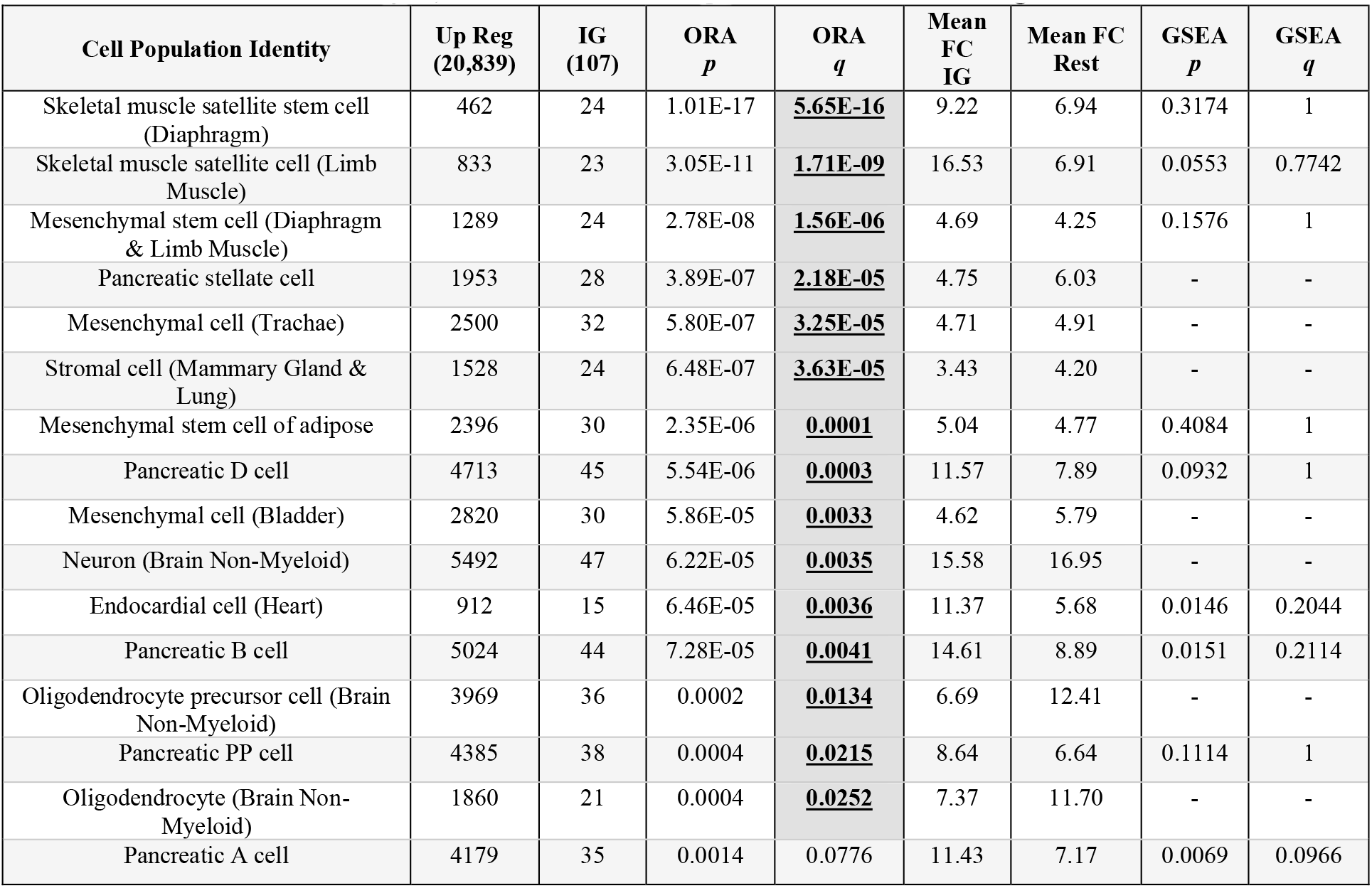
Imprinted gene over-representation in Tabula Muris adult cell populations (Schaum et al., 2018). Identity – Cell identities for the cells used in analysis; All other column descriptions can be found in the legend of Table 1.

### Imprinted genes are over-represented in embryonic and neonatal tissues vs. adult and demonstrate gene set enrichment in placental trophoblast progenitors

As the MCA adult tissues were sequenced alongside a selection of embryonic (e14.5) and neonatal tissues. The tissue analysis (Table 3, Supplemental Table S4) revealed a distinct bias for overrepresentation of imprinted genes in embryonic (5/9) and neonatal (5/6) tissues over adult tissues (1/18); the pancreas being the only adult tissue to be considered over-represented for imprinted genes when compared to the embryo and neonate. Both the placenta and embryonic mesenchyme were over-represented for imprinted genes.

**Table 3.**
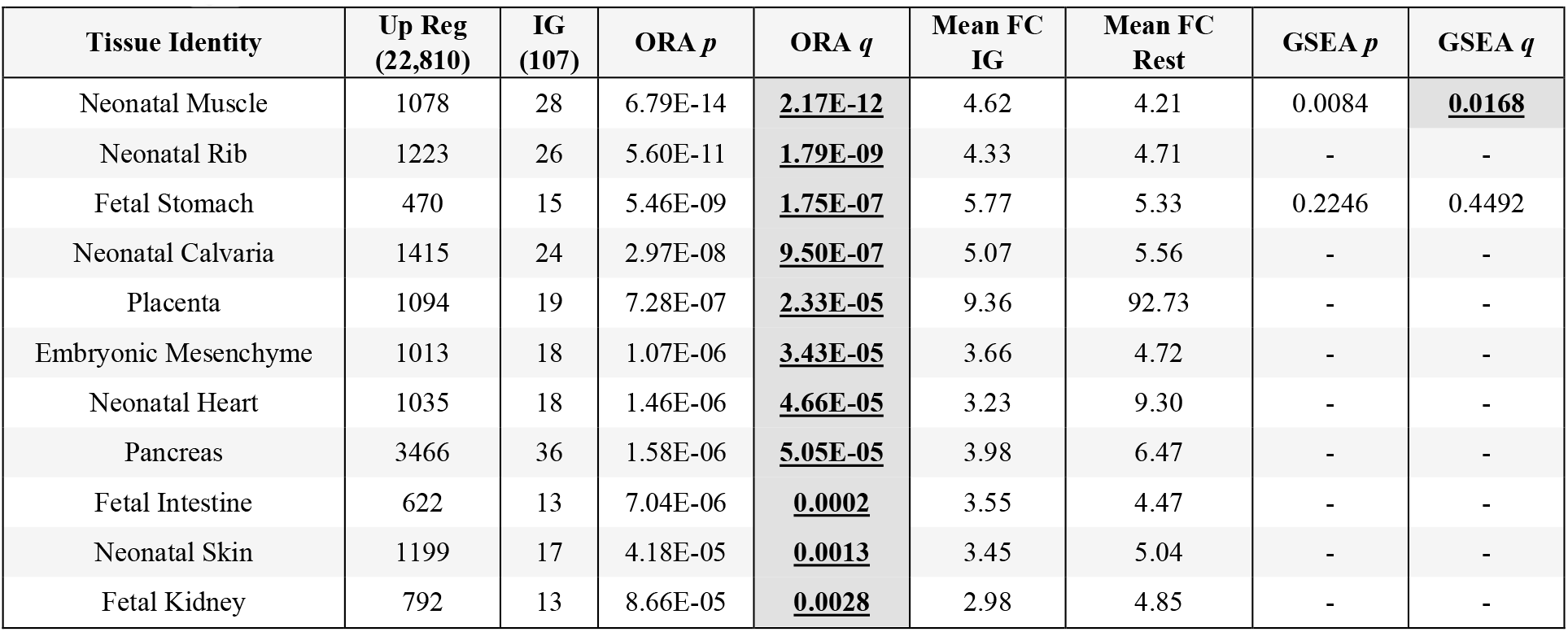
Imprinted gene enrichment in tissues derived from adult, foetal, embryonic and stem-cell derived cells in the MCA (Han et al., 2018). Identity – Tissue identities for the cells used in analysis; All other column descriptions can be found in the legend of Table 1.

| Tissue Identity | Up Reg (22,810) | IG (107) | ORA *p* | ORA *q* | Mean FC IG | Mean FC Rest | GSEA *p* | GSEA *q* |
| --- | --- | --- | --- | --- | --- | --- | --- | --- |
| Neonatal Muscle | 1078 | 28 | 6.79E-14 | **2.17E-12** | 4.62 | 4.21 | 0.0084 | **0.0168** |
| Neonatal Rib | 1223 | 26 | 5.60E-11 | **1.79E-09** | 4.33 | 4.71 | - | - |
| Fetal Stomach | 470 | 15 | 5.46E-09 | **1.75E-07** | 5.77 | 5.33 | 0.2246 | 0.4492 |
| Neonatal Calvaria | 1415 | 24 | 2.97E-08 | **9.50E-07** | 5.07 | 5.56 | - | - |
| Placenta | 1094 | 19 | 7.28E-07 | **2.33E-05** | 9.36 | 92.73 | - | - |
| Embryonic Mesenchyme | 1013 | 18 | 1.07E-06 | **3.43E-05** | 3.66 | 4.72 | - | - |
| Neonatal Heart | 1035 | 18 | 1.46E-06 | **4.66E-05** | 3.23 | 9.30 | - | - |
| Pancreas | 3466 | 36 | 1.58E-06 | **5.05E-05** | 3.98 | 6.47 | - | - |
| Fetal Intestine | 622 | 13 | 7.04E-06 | **0.0002** | 3.55 | 4.47 | - | - |
| Neonatal Skin | 1199 | 17 | 4.18E-05 | **0.0013** | 3.45 | 5.04 | - | - |
| Fetal Kidney | 792 | 13 | 8.66E-05 | **0.0028** | 2.98 | 4.85 | - | - |

Cells from this large comparison were then recategorized into 67 tissue/timepoint-spanning cell types (Table 4, Supplemental Table 5). Over-representation was seen principally in the mesenchymal stem cell derived cells (osteoblast, chondrocyte, myocyte, muscle cell, cartilage) spread across neonatal and adult tissue. In addition, endocrine cells mainly derived from the pancreas were over-represented. Only the trophoblast progenitor cells of the placenta showed a significant GSEA for imprinted genes (Figure 2).

**Table 4.**
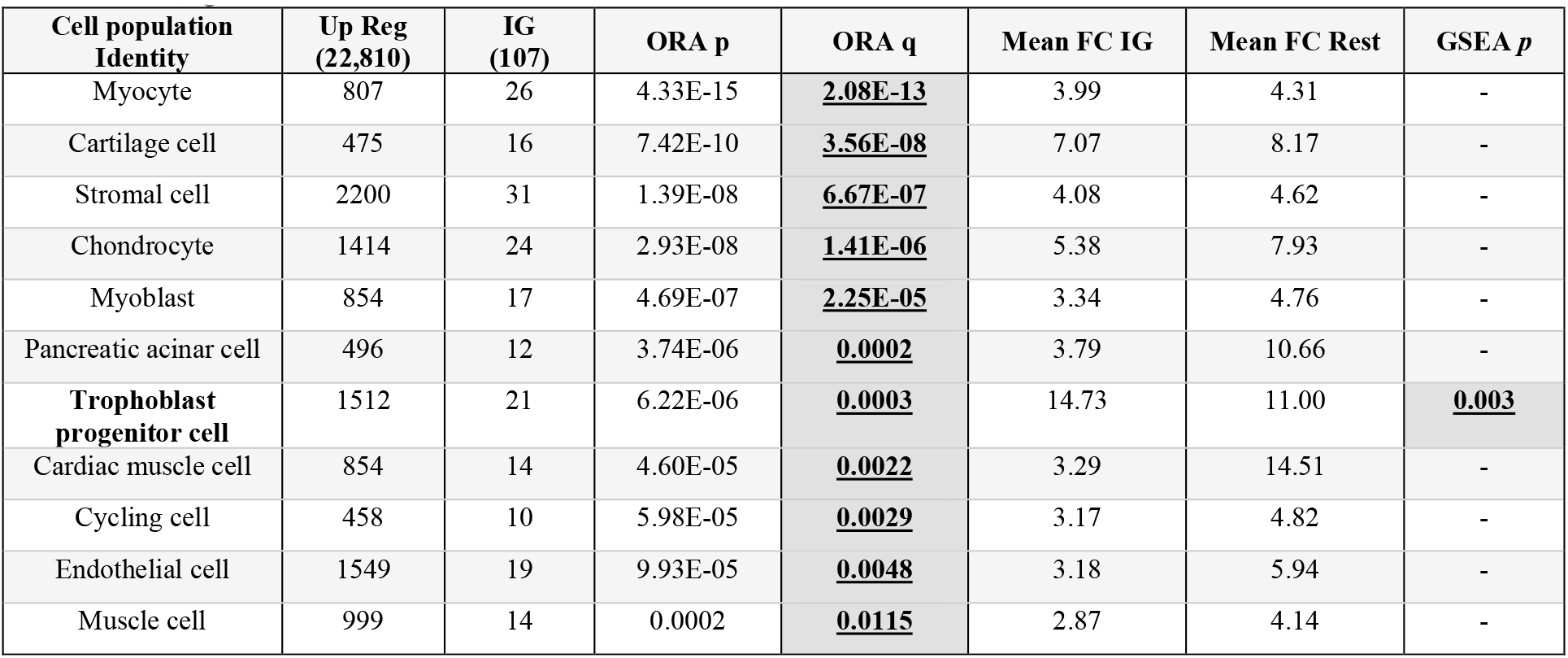
Imprinted gene enrichment in global cell types across adult, foetal, embryonic and stem-cell derived tissues in the MCA (Han et al., 2018). Identity – Cell identities for the cells used in analysis; All other column descriptions can be found in the legend of Table 1.

**Figure 2.**
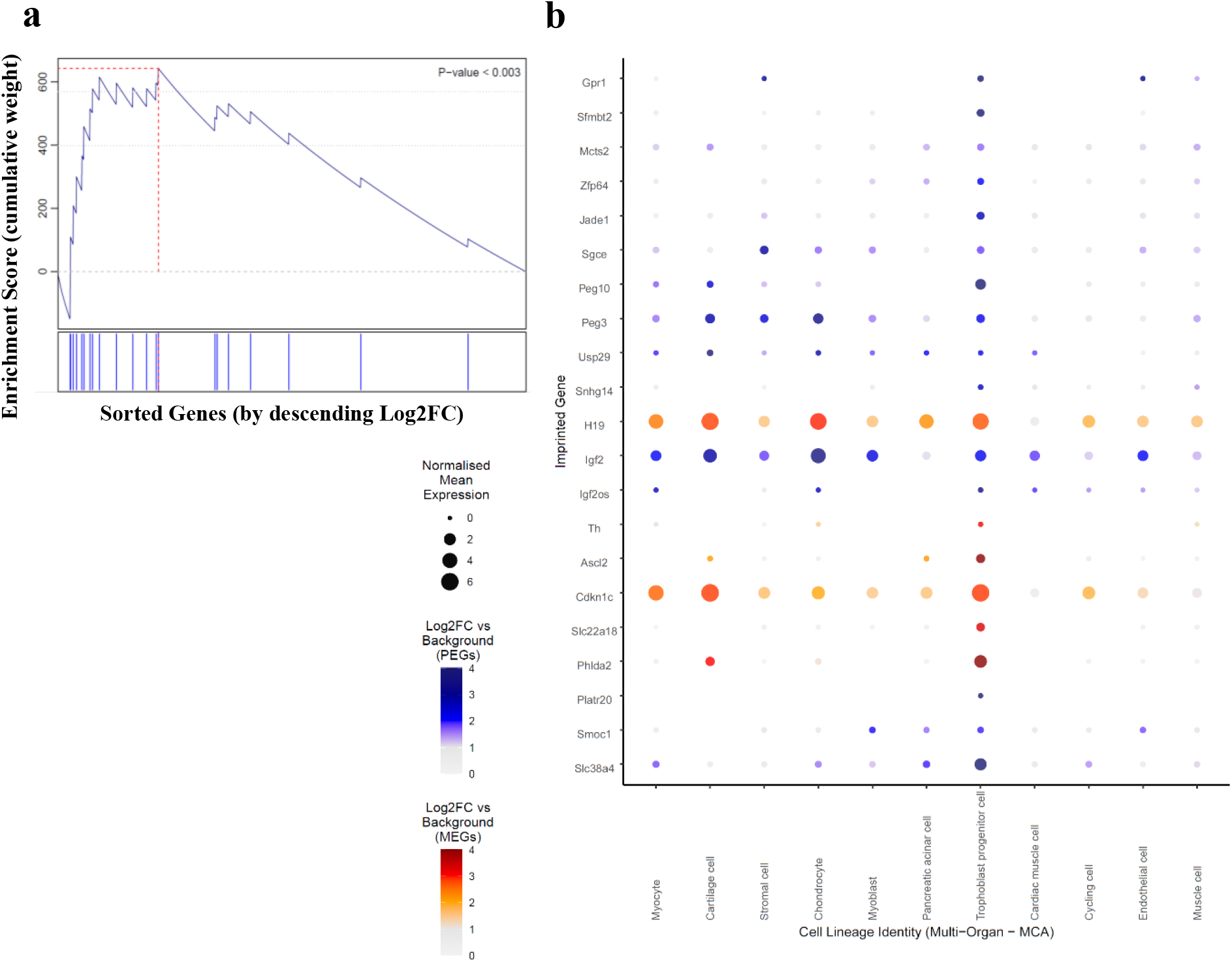
Gene Set Enrichment Analysis of Imprinted Genes for genes upregulated in Progenitor Trophoblast cells. (A) GSEA graph - In the analysis, genes are sorted by strength by which they mark this neuronal cluster (sorted by Log2FC values) indicated by the bar (bottom). The genes are arrayed left (strongest marker) to right and blue lines mark where imprinted genes fall on this array. The vertical axis indicates an accumulating weight, progressing from left to right and increasing or decreasing depending on whether the next gene is an imprinted gene or not. The p-value represents the probability of observing the maximum value of the score (red dashed line) if the imprinted genes are distributed randomly along the horizontal axis. (B) Dot plot of imprinted genes upregulated in the Trophoblast Progenitor Cells plotted across all imprinted gene over-represented cell types in the MCA. Size of points represented absolute mean expression; colour represented the size of the Log2FC value for that cell type (e.g., trophoblast progenitor cells) vs. all other cells. MEGs and PEGs possess distinct colour scales.

### Imprinted genes are over-represented in pancreatic endocrine cell types

By whole cells analysis of the MCA imprinted genes were over-represented in the pancreatic endocrine cells and the beta cells alongside the three stromal cell subpopulations (Table 1). When analysing the pancreas alone (Supplemental Table 6a), the stromal cells were the only cell type with an over-representation of the number of imprinted genes. However, although beta cells did not show over representation, imprinted genes were expressed at 3.9x the mean FC with dramatically higher levels of expression.

The TM identified a similar selection of pancreatic cells. Within the tissue-wide analysis (Table 2) we identified over-representation of imprinted genes in three of the four types of endocrine islet cells in the pancreas (Beta, Delta and PP, but not Alpha) alongside the top hit, the stellate cells - a multifunctional cell in the endocrine and exocrine pancreas that express collagen and fibronectin and promotes fibrosis when activated (Zhou et al., 2019). When analysing the pancreas cells alone (Supplemental Table 6b) over-representation was seen in the delta cells (q = 0.004) and the stellate cells (q = 0.033).

To provide another independent pancreatic cell analysis, we analysed the pancreatic data from Baron et al. (2016) (Table 5). Again, consistent with the TM and MCA, imprinted genes were over-represented in the endocrine cell populations of the pancreas (although this time all 4 types), with the δ and α cells as the top hits (Figure 3A). Figure 3B & 3C present dot plots of the imprinted genes upregulated in the cell populations within the pancreatic datasets including all the islets cells (TM and Baron et al. (2016)).

**Table 5.**
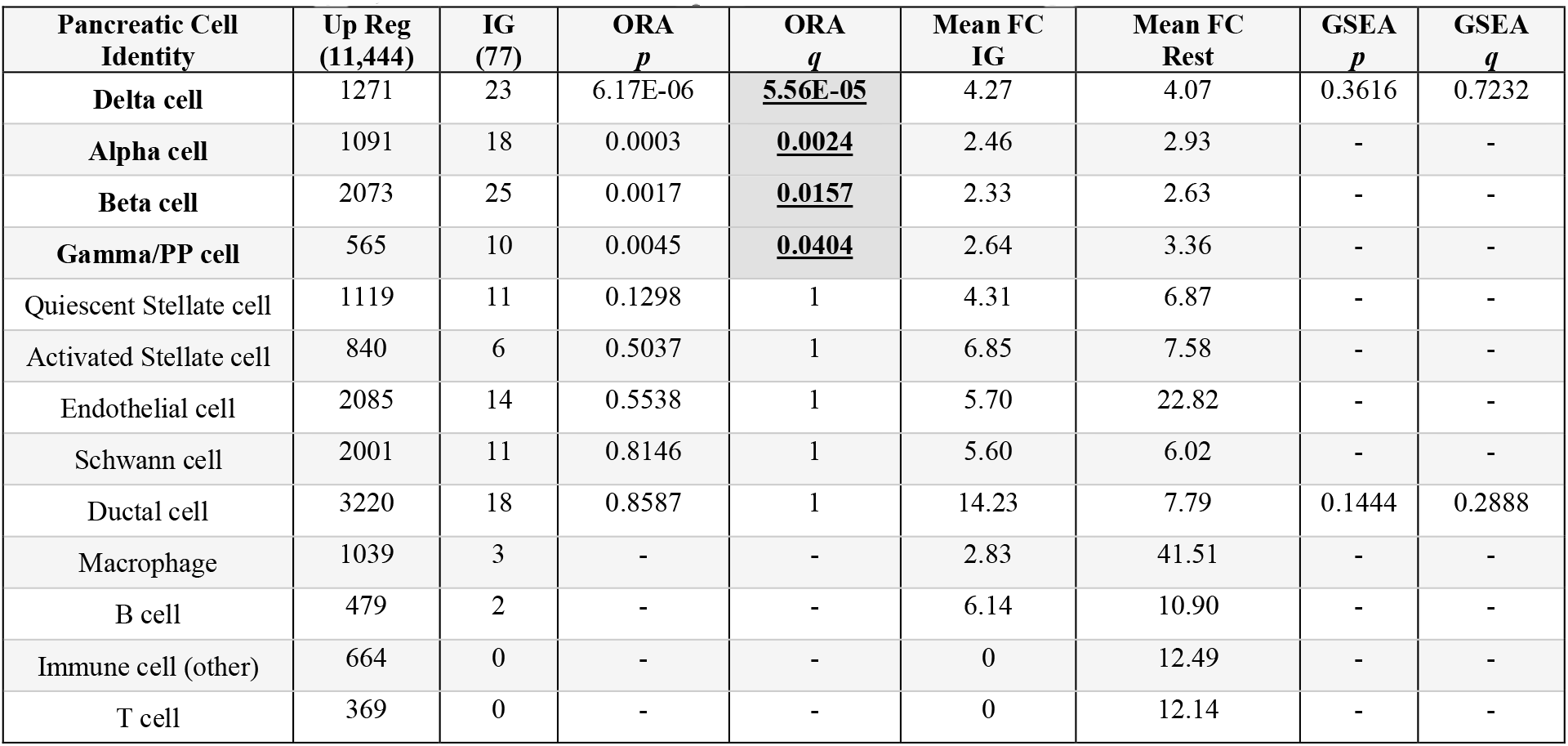
Imprinted gene over-representation in Pancreatic adult cell populations (Baron et al., 2016). Identity Cell identities for the cells used in analysis; All other column descriptions can be found in the legend of Table 1.

| Pancreatic Cell Identity | Up Reg (11,444) | IG (77) | ORA $p$ | ORA $q$ | Mean FC IG | Mean FC Rest | GSEA $p$ | GSEA $q$ |
| --- | --- | --- | --- | --- | --- | --- | --- | --- |
| Delta cell | 1271 | 23 | 6.17E-06 | <u>5.56E-05</u> | 4.27 | 4.07 | 0.3616 | 0.7232 |
| Alpha cell | 1091 | 18 | 0.0003 | <u>0.0024</u> | 2.46 | 2.93 | - | - |
| Beta cell | 2073 | 25 | 0.0017 | <u>0.0157</u> | 2.33 | 2.63 | - | - |
| Gamma/PP cell | 565 | 10 | 0.0045 | <u>0.0404</u> | 2.64 | 3.36 | - | - |
| Quiescent Stellate cell | 1119 | 11 | 0.1298 | 1 | 4.31 | 6.87 | - | - |
| Activated Stellate cell | 840 | 6 | 0.5037 | 1 | 6.85 | 7.58 | - | - |
| Endothelial cell | 2085 | 14 | 0.5538 | 1 | 5.70 | 22.82 | - | - |
| Schwann cell | 2001 | 11 | 0.8146 | 1 | 5.60 | 6.02 | - | - |
| Ductal cell | 3220 | 18 | 0.8587 | 1 | 14.23 | 7.79 | 0.1444 | 0.2888 |
| Macrophage | 1039 | 3 | - | - | 2.83 | 41.51 | - | - |
| B cell | 479 | 2 | - | - | 6.14 | 10.90 | - | - |
| Immune cell (other) | 664 | 0 | - | - | 0 | 12.49 | - | - |
| T cell | 369 | 0 | - | - | 0 | 12.14 | - | - |

**Figure 3.**
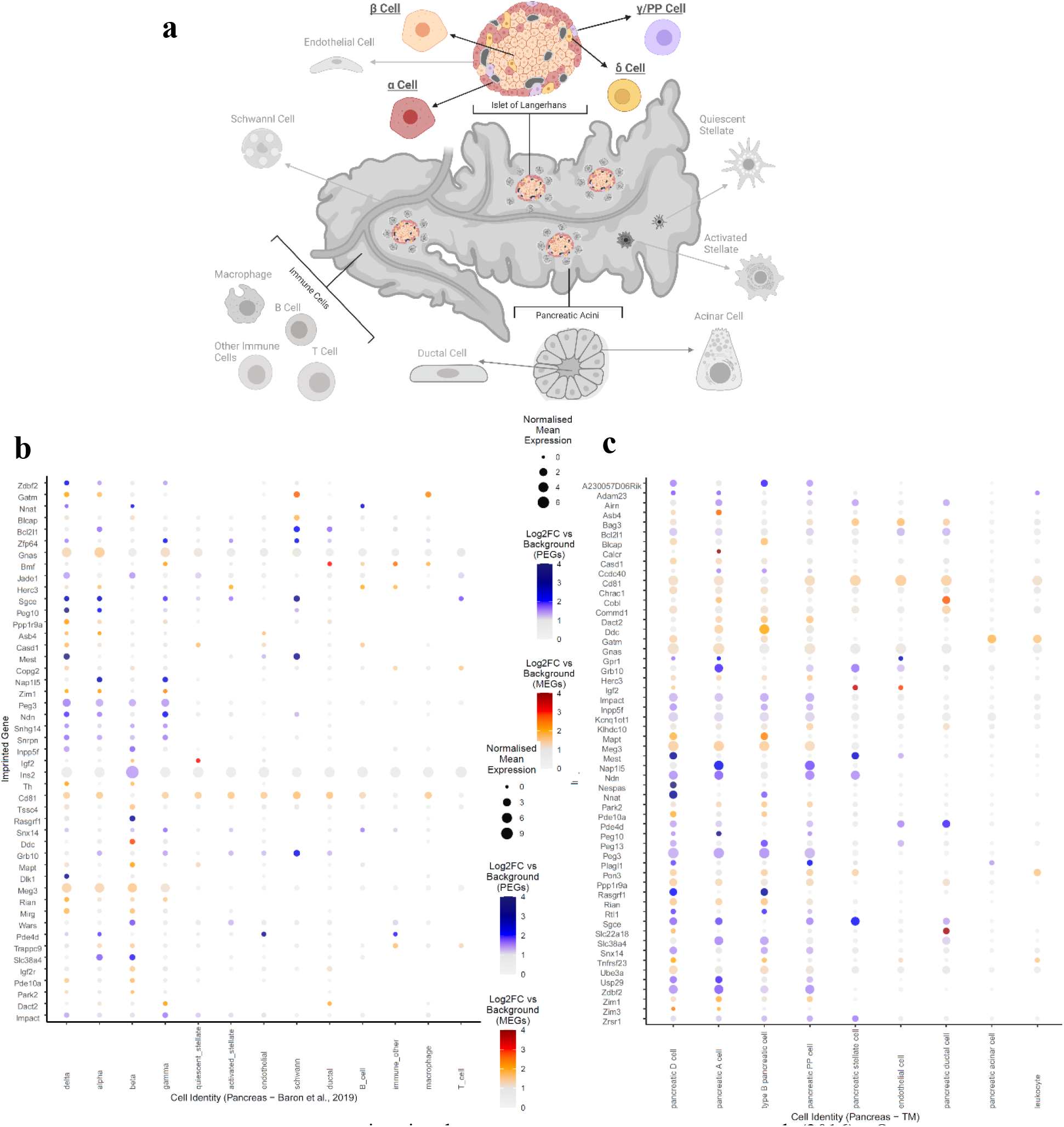
(A) Pancreatic cell types with imprinted gene over-representation in Baron et al. (2016) - Over-represented cell types are bold, underlined and not in greyscale. (B&C) Dot Plots showing imprinted gene expression across all cell types in Baron et al. (2016) (B) and Tabula Muris (C) pancreatic datasets – Imprinted genes with upregulated expression in at least one endocrine cell were plotted for each dataset independently across all cell types. See legend of Figure 2 for how to interpret the plot.

### Imprinted genes are over-represented in mesenchymal bladder cells

We saw an over-represented bladder stromal cell population in the MCA (Table 1), potentially explaining why we found it as an over-represented tissue previously (Higgs et al., 2022). The authors of the MCA distinguished bladder stromal cells as expressing Bmp4 and Wnt2, both of which are markers of mesenchymal stromal cells (Pokrywczynska et al., 2019, Mysorekar et al., 2002) with a role in epithelial maintenance and renewal (Mysorekar et al., 2009). Convergently, the mesenchymal bladder cells were the over-represented population in the TM analysis (Table 2), suggesting this cell type is the reason why imprinted genes show enriched expression in the bladder.

### Imprinted genes are over-represented in Muscle satellite/stem cells

For muscle-based tissues, the MCA cell subpopulation analysis found imprinted gene overrepresentation in two muscle-based cell subpopulations (Table 1): a stromal cell population and a muscle progenitor population. In the TM analysis (Table 2), over-representation was seen in a variety of cells originating from the muscle tissues as the top hits – skeletal muscle satellite cells and mesenchymal stem cells, which mirrors the enrichment of muscle progenitor cells and the mesenchymal cells mirroring the stromal cells in the MCA.

The data from De Micheli et al. (2020) of skeletal muscle during repair were analysed as another independent dataset of muscle cells (Table 6). We saw significant GSEAs for imprinted genes in the stem cell and progenitor cell types (also known as the satellite cells) (Figure 4). Additional over-representation was seen in the fibro/adipogenic progenitor cells, also a type of mesenchymal stromal cell which support satellite cell differentiation (Biferali et al., 2019). This shows a very consistent picture with the MCA and TM analyses that the mesenchymal stem cell populations in the muscle are the key imprinted gene enriched population.

**Table 6.** Imprinted gene over-representation in Muscle cell populations (De Micheli et al., 2020). Identity – Cell identities for the cells used in analysis; All other column descriptions can be found in the legend of Table 1.

| Cell Population Identity | Up Reg<br>(17,072) | IG<br>(100) | ORA<br>*p* | ORA<br>*q* | Mean FC<br>IG | Mean FC<br>Rest | GSEA<br>*p* | GSEA<br>*q* |
| --- | --- | --- | --- | --- | --- | --- | --- | --- |
| FAPs | 8270 | 69 | 2.43E-05 | **0.0003** | 5.59 | 4.25 | 0.1473 | 0.4419 |
| Smooth Muscle Cells | 5761 | 45 | 0.0124 | 0.1492 | 5.11 | 3.73 | 0.3303 | 0.9909 |
| MuSCs and Progenitors | 8191 | 59 | 0.0173 | 0.2074 | 8.12 | 2.91 | 2.00E-04 | **6.00E-04** |
| Neural, Glial & Schwann Cells | 2182 | 20 | 0.0269 | 0.3225 | 5.60 | 26.73 | - | - |
| Mature Skeletal Muscle | 2221 | 16 | 0.2239 | 1 | 4.84 | 10.91 | - | - |
| Endothelial | 2888 | 20 | 0.2400 | 1 | 3.03 | 5.43 | - | - |
| Tenocytes | 2655 | 17 | 0.3853 | 1 | 2.52 | 7.33 | - | - |
| Monocytes, Macrophages & Platelets | 4754 | 17 | 0.9960 | 1 | 1.46 | 1.77 | - | - |
| Pro-inflammatory Macrophages | 2984 | 8 | 0.9980 | 1 | 1.32 | 1.92 | - | - |
| Anti-inflammatory Macrophages | 3686 | 10 | 0.9993 | 1 | 2.48 | 2.12 | - | - |
| B, T, NK Cells | 2590 | 5 | 0.9996 | 1 | 4.04 | 11.61 | - | - |
| Resident Macrophages APCs | 3580 | 8 | 0.9999 | 1 | 1.50 | 3.77 | - | - |

**Figure 4.**
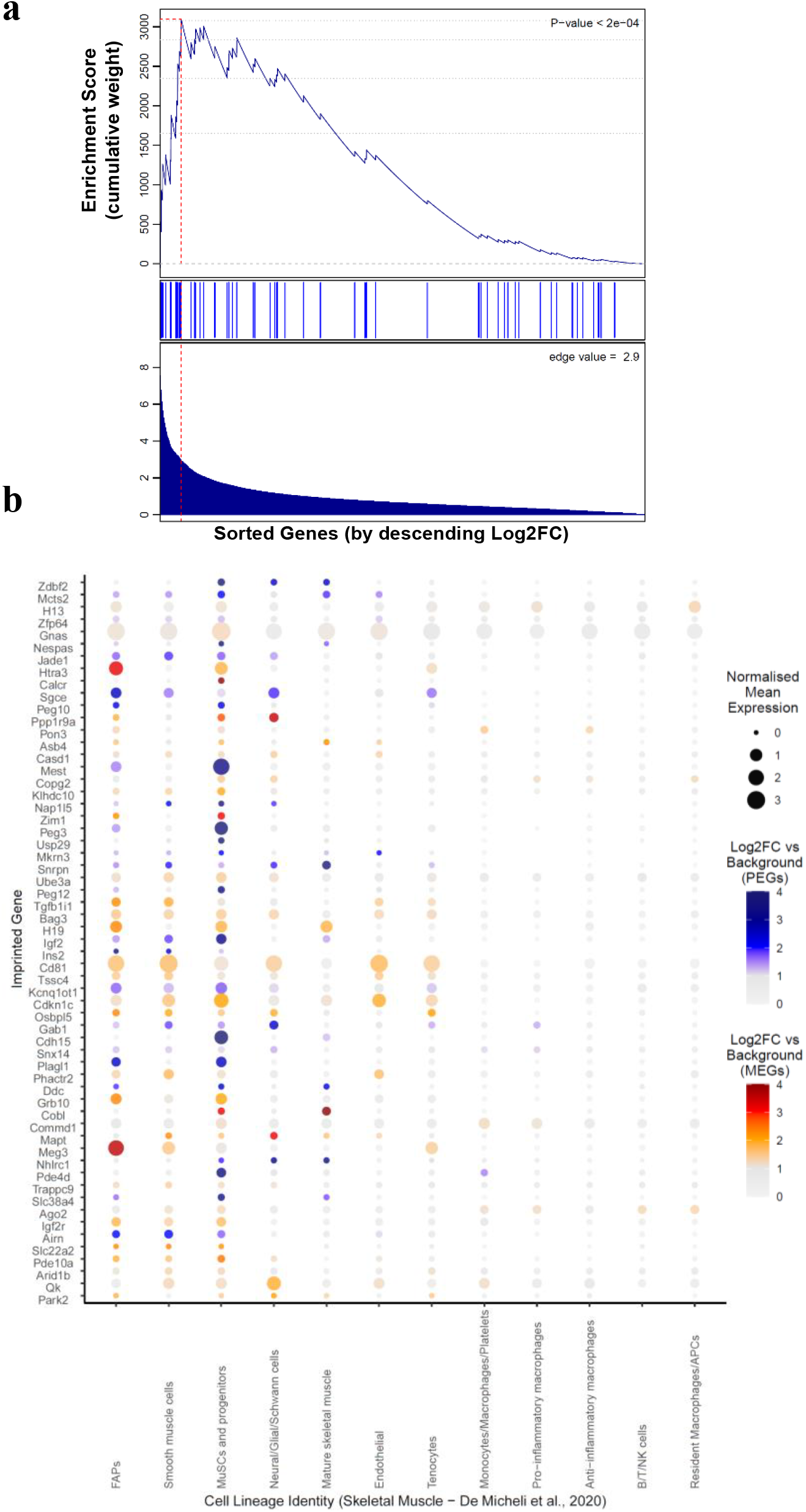
Gene Set Enrichment Analysis of Imprinted Genes for genes upregulated in MuSCs and Progenitors from De Micheli et al. (2020). (A) GSEA for imprinted genes upregulated in the ‘MuSCs and progenitors’ cell type in the Skeletal muscle dataset. See legend of Figure 2 for a description of how to interpret the plot. (B) Dot plot of imprinted genes upregulated in the ‘‘MuSCs and progenitors’ cell type plotted across all identified cell types in the De Micheli et al. (2020) skeletal muscle dataset. See legend of Figure 2 for a description of how to interpret the plot.

### Imprinted genes are over-represented in Mammary Gland stem cell populations

The mammary gland was not an enriched tissue in either the MCA or TM multi-tissue analysis (Higgs et al., 2022), but the MCA cell subtype analysis saw populations of mammary stromal cell over-represented, and the stromal cells derived from the mammary gland were also enriched in the TM. Both datasets distinguished these stromal cells by expression of matrix metallopeptidases (Mmp). Mammary Stroma consists of a variety of cells including adipocytes, fibroblasts and ECM. The structural ECM is a large part of the stroma and is believed to be an important mammary stem cell niche (Wiseman and Werb, 2002, Khokha and Werb, 2011).

Recently, Xu et al. (2020) reported high expression of imprinted genes in a stem-cell type in the epithelial component of the mammary gland in the Bach et al. (2017) sequencing of the epithelial cells and over-representation analysis of this scRNA-seq dataset of mammary epithelial cells shows imprinted genes are over-represented in the basal compartment of the mammary epithelium, but the top hit was the stem (Procr+) cells as Xu et al. (2020) reported (Table 7).

**Table 7.** Imprinted gene over-representation in Mammary Gland Epithelial cells (Bach et al., 2017) Identity – Cell identities for the cells used in analysis; All other column descriptions can be found in the legend of Table 1.

| Cell Population Identity | Up Reg<br>(16,415) | IG<br>(96) | ORA<br>p | ORA<br>q | Mean FC<br>IG | Mean FC<br>Rest | GSEA<br>p |
| --- | --- | --- | --- | --- | --- | --- | --- |
| Mammary Gland Stem (*Procr+*) cells | 1267 | 21 | 1.06E-05 | **8.46E-05** | 25.11 | 30.79 | - |
| Basal cells | 1446 | 18 | 0.0016 | **0.0132** | 6.88 | 6.22 | 0.2045 |
| Luminal hormone sensing differentiated | 2120 | 18 | 0.0649 | 0.5194 | 5.39 | 6.63 | - |
| Myoepithelial cells | 2343 | 15 | 0.3952 | 1 | 6.42 | 8.48 | - |
| Luminal hormone sensing progenitors | 1217 | 8 | 0.4194 | 1 | 2.40 | 2.66 | - |
| Luminal progenitor cells | 982 | 5 | 0.6876 | 1 | 3.74 | 4.13 | - |
| Luminal alveolar cells differentiated | 2228 | 11 | 0.7705 | 1 | 5.80 | 8.05 | - |
| Luminal alveolar cells progenitor | 1113 | 5 | 0.7879 | 1 | 3.02 | 4.88 | - |

We next asked whether imprinted genes were over-represented at different stages of the mammary gland cycle (Virgin, Gestation, Lactation and Involution) (Hanin and Ferguson-Smith, 2020). The MCA dataset sequenced mammary gland tissue at these different stages of the pregnancy cycle and hence allowed us to analyse whether imprinted genes were over-represented in the mammary gland cells at a particular life period. Although the largest number of imprinted genes were seen during involution and pregnancy, no specific time point was over-represented for imprinted genes (Supplemental Table S7). When analysing all cell types at each pregnancy timepoint, stromal cells were the only over-represented cell for each timepoint (Supplemental Table 8). A significant GSEA was only seen for pregnancy mammary stromal cells. Analysis of the epithelial cells only (Bach et al., 2017) at the four mammary periods saw no significant over-representation for any time-point (Supplemental Table 9).

## DISCUSSION

Here, we applied a systems biology approach to identify tissues and cell subpopulations enriched for imprinted genes as evidence of their convergent functional relevance. We began our analysis by looking for enrichment amongst all adult cell subpopulations taken from the MCA and TM mouse organ single cell compendiums. We saw a generalised enrichment in the mesenchymal stromal cells, a population of cells with known cell renewal capacities, and most likely the principal cell-type responsible for imprinted gene enrichment seen in the bladder, muscle tissues and the stromal cell-types in the mammary gland too. Cell-types that were enriched for imprinted genes outside these stromal/stem cell-types were principally the endocrine pancreas cells alongside the pituitary trophic cells, and the neurons identified previously (Higgs et al., 2022). These ‘expression hotspots’ point to ‘functional hotspots’. Consequently, our findings highlight a key role for imprinted genes in stemness and tissue maintenance, within parenchymal cells, and in neuro-endocrine regulation.

The endocrine cell subpopulations of the islets of Langerhans in the pancreas produce specific hormones, namely glucagon from alpha cells, insulin from beta cells, somatostatin from delta cells and ghrelin from PP cells. These cells interact with each other in paracrine feedback to regulate the metabolism of glucose. Imprinted genes are known to have a role in metabolism, regulation of insulin and of blood glucose levels (Millership et al., 2019) and some genes have been associated with the endocrine pancreas (Stefan et al., 2011). Specific regulation of the beta pancreatic cells has been demonstrated for imprinted genes such as Rasgrf1(de Mora et al., 2003) and Nnat (Millership et al., 2018), two genes upregulated in this cell type in this study. However, our analysis here suggests that imprinted genes may have a role in the islet cells collectively (alpha and delta included), since insulin and glucose regulation can also be performed indirectly by influencing the hormonal output of the alpha and delta cells. As seen from Figure 3, the relevant imprinted genes are not expressed continuously through the endocrine cell populations but for the most part display a cell-type specific enrichment, which suggests direct regulation of one hormone output in the pancreas, having paracrine effects on the others, for individual imprinted genes.

Mesenchymal/stromal cells were the other major cell type identified in the adult tissue compendiums and the specific analyses of the muscle tissue and mammary gland. These cells represent a variety of cell types, specifically the fibroblasts, extracellular matrix (ECM) and mesenchymal stem cells, which make up the connective support tissue around parenchymal cells. Imprinted genes have been widely reported to influence cell-cycle length, proliferation and tumor formation (Hernandez et al., 2003, Al Adhami et al., 2015, Plasschaert and Bartolomei, 2014) and to be expressed in adult stem cells and subsequently repressed to allow self-renewal (Berg et al., 2011, Besson et al., 2011, Godini et al., 2019, Zacharek et al., 2011). Hence, we were not surprised to find enrichment in the mesenchymal stem cells/satellite cells in muscle tissues, a cell type crucial for muscle regeneration which is a physiological process in which imprinted genes have been associated (Moresi et al., 2015, Yan et al., 2003). Furthermore, the satellite cells have been of direct interest for genes such as H19 (Martinet et al., 2016) and Peg3 (Correra et al., 2018). However, we suspect the ECM component of the stromal cell population is also contributing to this enrichment. Within the imprinted gene networks, ECM genes were heavily co-expressed alongside imprinted genes, becoming incorporated into the network (Varrault et al., 2006) and Al Adhami et al. (2015) suggested that imprinted genes can regulate cell proliferation and cell cycle exit, not just directly through the impacting the cell machinery as shown for Cdkn1c (Matsuoka et al., 1995) but indirectly through modulation of the ECM composition to impact proliferation-quiescence-differentiation (Brizzi et al., 2012). An enrichment of imprinted genes amongst the stromal cells supports this idea and also recent suggestions that the imprinted genes have a role in stemness, maintenance, cell differentiation and lineage determination in the mammary gland (Xu et al., 2020), Finally, when analysing global cell types from across the developmental spectrum in the MCA (fetal, neonatal and adult) the mesenchymal cell lineage were the consistent top hits, particularly during fetal and neonatal development. Cell types such as the endocrine pancreas were still represented at this level of analysis but the only cell type identified by the more conservative GSEA were the trophoblast progenitor cells of the placenta. The placenta is a key site of action for imprinted genes (Coan et al., 2005, Fowden et al., 2006, John, 2017, Peters, 2014, Haig, 1996). Recent work has suggested that expression of these genes during the cell differentiation of trophoblast progenitor cells has important functional outcomes impacting the endocrine capacity of the placenta, with knock-on effects on maternal behaviour (Creeth et al., 2019). A large number of canonical imprinted genes have demonstrated monoallelic expression in the trophoblast stem cells (Calabrese et al., 2015) and manipulation of imprinted genes has been shown to cause loss or gain of specific placental lineages e.g., Phlda2 (Tunster et al., 2016a), Cdkn1c (Tunster et al., 2011), Peg3 (Tunster et al., 2018), Peg10 (Ono et al., 2003), Igf2 (Esquiliano et al., 2009) and Ascl2 (Tunster et al., 2016b). It is therefore reassuring and valuable to see that, although most imprinted genes are not expressed in this cell type (hence not over-represented), the ones that are expressed are amongst the top expressed genes, so that they alone create a significant GSEA. The genes already shown to have placental lineage phenotypes all demonstrated dramatic upregulation in the trophoblast progenitor cells as well as a few other genes that have not been studied which represent important candidates for future placental analysis.

The limitations of the current work are as previously reported (Higgs et al., 2022). Briefly, we were bound by data and intra-dataset cellular comparisons that were available to us and not all avenues or intricate cell types could be considered. Our approach using ORA and GSEA do not provide an exhaustive list of expression sites, but the most significant sites of disproportionally large numbers of upregulated imprinted genes. Finally, as before, we did not assess parent-of-origin expression for the 119 imprinted genes we included in the analysis in all cell subpopulations. We have, however, restricted our gene selection to genes with reliable imprinting status (the canonical imprinted genes and genes with more than one demonstration of a POE) when looking for enrichment. Although this does not replace validating the imprinting status of all 119 in all the cell subpopulations examined, it does provide justification for looking at imprinted gene over-representation.

In summary, this extended scRNA-seq analysis asked whether imprinted genes, as a group, were disproportionately represented in developing and adult mouse cell types. We discovered that imprinted genes show a body-wide enrichment pattern within mesenchymal/stromal cells, representing the cell populations responsible for tissue maintenance and cell replacement. Of further interest were the non-stem cell populations found enriched for imprinted genes, namely the endocrine pancreas cells and progenitors of the placental endocrine lineages. In summary, our investigations indicate that imprinted genes, when considered as a set, are principally upregulated in the cell maintenance/differentiation related mesenchymal/stromal cells and the neuro-hormonal system of the mouse body.

## METHODS

### Data Processing

Five unique datasets were analysed across the two levels of analysis (see Fig.1) and analyses were conducted on each dataset independently. We aimed to be unbiased by using all the datasets that sequenced the tissues identified by the Level 1 Multi-Organ analysis in Higgs et al. (2022). All sequencing data were acquired through publicly available resources and each dataset was filtered and normalised according to the original published procedure. Table 8 details the basic parameters of each dataset. Once processed, each dataset was run through the same basic workflow (see below and Figure 5), with minor adjustments laid out for each dataset detailed in the Supplemental Methods.

**Table 8.** Dataset specific sequencing and processing information for all datasets analysed. Datasets are organised by level of analysis and compared for single-cell sequencing protocol, Animal and Tissue processing, Cell Quality Filters used, No. of Cells in final dataset and Data Normalisation procedure followed.

| Dataset | Protocol | Animal/Tissue | Cell Filter | Filtered cell no. | Normalisation |
| --- | --- | --- | --- | --- | --- |
| *Mouse Cell Atlas (Han et al., 2018)* | Microwell-seq | C57BL/6J, M+F, 6-10 weeks, E14.5 and neonatal | 1500 highest quality cells per tissue | 61,637 | 100,000 transcripts log transformed |
| *Tabula Muris (Schaum et al., 2018)* | Smart-seq2 (FACS) | C57BL/6J 7 mice (4F/3M), 10-15 weeks, virgin | reads>50,000<br>genes>500 | 44,879 | ln(CPM+1) |
| *Pancreas (Baron et al., 2016)* | inDrop | C57BL/6 and ICR mice, M+F, 5 mice | Reads > 1,000<br>Genes > 750 | 1,886 | Log(TPM+1) |
| *Skeletal Muscle (De Micheli et al., 2020)* | 10X Chromium | C57BL/6J, M+F, 3-7 months of age, tibialis anterior muscles | UMI > 1000<br>Genes > 200 | 34,438 | By total UMI per cell, log transform |
| *Mammary Gland (Bach et al., 2017)* | 10X Chromium | C57BL/6J, F, ~8 week at different lactation stages (e14.5, p6, p11 and NP) | UMI > 1000<br>genes > 500<br>mito < 0.05 | 23,184 | Size Factors<br>Scran,<br>log2 transform |

**Figure 5.**
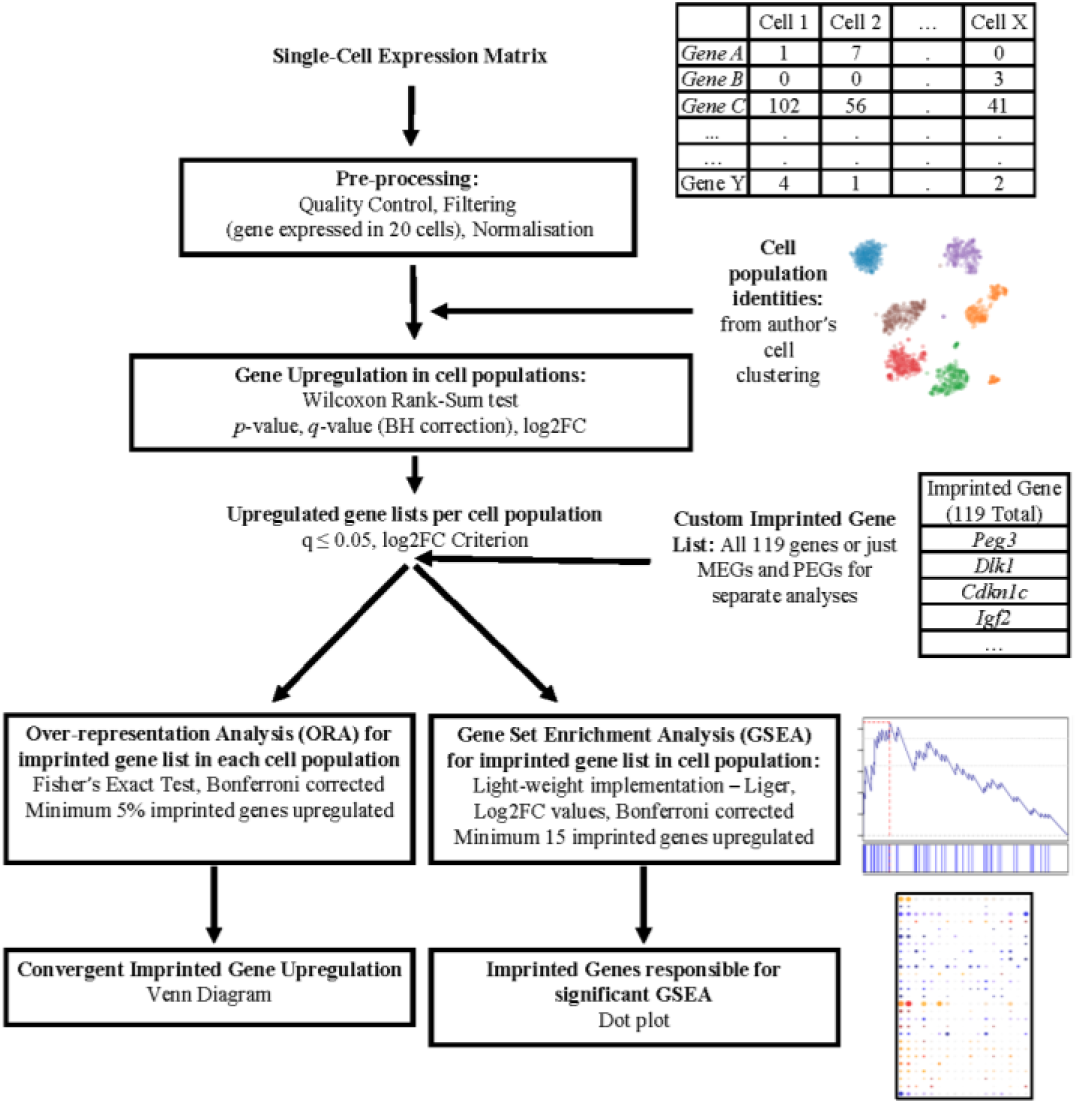
Basic workflow schematic. Single Cell Expression Matrices were acquired through publicly available depositories. Data were processed according to the author’s original specifications and all genes were required to be expressed in 20 or more cells. Cell population identities were acquired from the author’s original clustering. Positive differential gene expression was calculated via Wilcoxon Rank-Sum Test. Upregulated genes were considered as those with q ≤ 0.05 and a Log2FC ≥ 1 for analysis levels 1 and this criterion was relaxed to Log2FC > 0 for level 2. Our imprinted gene list was used to filter upregulated genes and two different enrichment analyses were carried out, over-representation analysis via Fisher’s Exact Test and Gene Set Enrichment Analysis via Liger algorithm (https://github.com/JEFworks/liger). Venn diagrams and dot plots were utilised for visualisation.

Due to the high variability in sequencing technology, mouse strain, sex and age, and processing pipeline, we have avoided doing analysis on combined datasets. Rather we chose to perform our analyses independently for each dataset and look for convergent patterns of imprinted gene enrichment between datasets on similar tissues. As with any single-cell experiment, the identification of upregulation or over-representation of genes in a cell-type depends heavily on which other cells are included in the analysis to make up the ‘background’. Analysing separate datasets (with overlapping cell-types alongside distinct ones) and looking for convergent patterns of enrichment is one way of counteracting this limitation.

### Basic Workflow

A detailed account of our workflow approach to identify imprinted gene enrichment (see Figure 5) can be found in Higgs et al. (2022). In brief, data were downloaded in the available form provided by the original authors (either raw or processed) and, where necessary, were processed (filtered, batch-corrected and normalized) to match the author’s original procedure. A consistent filter, to remove all genes expressed in fewer than 20 cells, was applied to remove genes unlikely to play a functional role due to being sparsely expressed. Cell identities were supplied using the outcome of cell clustering carried out by the original authors, so that each cell included in the analysis had a cell-type or tissue-type identity. Cells were used from mice of both sexes when provided.

Positive differential expression (upregulation) was carried out using a one-sided Wilcoxon rank-sum test performed independently for each gene and for each identity groups vs. all other groups and horizontal Benjamini-Hochberg correction was applied to all p values. Following the standard of our previous work, for all analyses of cells from a specific tissue (Level 2), significant positive differential expression was considered as q <= 0.05 and a positive Log2 fold change (Log2FC) compared to background. When cells were compared across tissues from the whole organism (Level 1), the criteria for upregulated genes included demonstrating a Log2FC value of 1 or larger (i.e., 2-fold-change or larger), which selects for genes with distinctive upregulation in these, akin to a marker gene for that cell type.

Over-Representation Analysis (ORA) of imprinted genes in cell identity groups using one-way Fisher’s Exact Tests (‘fisher.test’ function in R core package ‘stats v3.6.2’) was the first metric of enrichment. A baseline of ≥5% of the imprinted genes upregulated in an identity group was applied to avoid over-representation analysis on minimal genes. Subsequent p-values for all identity groups with 5% or more imprinted genes were corrected using a Bonferroni correction. This provided a measure of whether imprinted genes are expressed as a set above expectation in particular identity groups. Gene-Set Enrichment Analysis (GSEA) of imprinted genes in cell identity groups was also conducted using a publicly available, light-weight implementation of the GSEA algorithm in R (Subramanian et al., 2005) (https://github.com/JEFworks/liger). This test is more conservative, asking if imprinted genes are enriched among the strong markers of a cell identity group (those with the highest log fold change - Log2FC). Groups analysed were selected as having an minimum number of upregulated imprinted genes to measure enrichment for (minimum of 15 as suggested by the GSEA user guide (https://www.gsea-msigdb.org/gsea/doc/GSEAUserGuideFrame.html) and having an average fold change of the upregulated imprinted genes greater than the average fold change of the rest of the upregulated genes for that tissue. Again, multiple p values generated from GSEA were corrected using a Bonferroni correction. If no cell populations met these criteria, GSEA was not run and were not included in the results.

To further elucidate the genes responsible for significant GSEA’s, dot plots of the imprinted genes upregulated in that identity group were plotted across all identity groups with absolute expression and Log2FC mapped to size and colour of the dots, respectively. Dot plots or Venn diagrams were also reported to demonstrate the overlap between individual imprinted genes contributing to the over-representation of a cell identity found in multiple datasets for the same tissue.

Graphical representations and statistical analyses were conducted using R 3.6.2 (Team, 2019) in RStudio (Team, 2015). Diagrams in Figures 1, 3 & 5 were created with BioRender.com.

## DATA ACCESS

The datasets analysed during the current study were acquired from publicly available resources and are available in the following GEO repositories, Mouse Cell Atlas – GSE108097, Tabula Muris – GSE109774, Pancreas - GSE84133, Muscle Tissue - GSE143437, Mammary Epithelium - GSE106273. All analysis data generated in this experiment is provided as Supplemental Data and in the following Open Science Framework repository (https://osf.io/jx7kr/). Custom R scripts to analyse each dataset are provided as Supplemental Code and are available at https://github.com/MJHiggs/IG-Single-Cell-Enrichment.

## ACKNOWELDGEMENTS

This work was supported by a Wellcome Trust PhD studentship (220090/Z/20/Z). We would like to thank all the research groups that carried out the single-cell RNA sequencing that made this study possible and to particularly acknowledge Dr. Karsten Bach, Dr. Leonard Cheung and Dr. Peng Hu for help accessing the cell metadata for their associated studies. Author contributions: M.Higgs performed bioinformatic analysis, with input from M.Hill; M.Higgs., M.Hill, and A.Isles contributed to project design, data interpretation, and wrote the manuscript; all co-authors reviewed and edited the manuscript.

