## Supplemental Materials for "Tissue-wide scRNA-seq analysis reveals enrichment of imprinted genes in stem and endocrine cell-types in mice"

### Supplemental Methods

#### Specific Dataset information and workflow specifics

Mouse cell atlas (adult and developing mouse – 43 tissues)  (Han et al., 2018)

The Mouse Cell Atlas (MCA) involved the collection and sequencing of 50+ tissues in an attempt to distinguish genetic signatures of tissues across a mouse's life, of which 43 passed quality control measures. These tissues included: 22 adult tissues (6-10 weeks old), 11 fetal tissues (E14.5), 6 neonatal tissues and 4 stem cell derived tissues. An Expression matrix ('MCA_Figure2-batch-removed.txt.tar.gz') of 60,000 cells of high quality (~1500 cells from the 43 tissues) was downloaded from Figshare from (https://figshare.com/articles/MCA_DGE_Data/5435866) alongside the cell metadata ('MCA_Figure2_Cell.Info.xlsx' for all 43 tissues and 'MCA_CellAssignments.csv' for adult tissues only). Data were then scaled to 100,000 reads and log normalised. Cells were filtered to remove cells without an annotation or cell annotations that had less than 5 cells. For the analysis on all 43 tissues, the workflow was run twice, once for all 43 tissue identities ('Tissue' column) and once again for the 63 major cell identities ('Cluster ID column') across the 43 tissues. For the adult tissue only, there were a total of 20,744 cells assigned to 292 unique specific cell identities from the adult mouse, annotated from the 'annotations' column. The core workflow was run once for all the adults cells and then again for the tissue types on interest from the previous whole tissue analysis, namely the pancreas, bladder and mammary gland. Genes were filtered to 20 cells for each analysis.

Tabula Muris (adult mouse – 20 tissues) (Schaum et al., 2018)

The Tabula Muris Consortium released the Tabula Muris soon after the MCA. The Tabula muris comprises 20 adult mouse organs with a partial overlap with the MCA but sequenced these tissues to a much greater depth and using two sequencing methods. FACs sorting allowed characterisation of fewer cells than drop-seq from the organ but at much higher sensitivity and depth and was completed for all 20 organs. This was completed using 3 female mice and 4 male mice, all 10-15 weeks old and on a C57BL/6JN background. The FACs dataset was selected for analysis as it contained the largest number of tissues and the only one to include the brain (broken down as myeloid brain cells (neurons, glia, endothelial etc) and non-myeloid brain cells (microglia and macrophages)). Clustering for cells in this dataset allowed cell identification at the Tissue and Cell Subtype level. All data were downloaded through Figshare as Robjects containing raw, processed data (https://figshare.com/articles/Robject_files_for_tissues_processed_by_Seurat/5821263/1. Cell metadata were acquired as annotation csv's from the respective technology's Figshare page (FACS - https://figshare.com/articles/Single-cell_RNA-seq_data_from_Smart-seq2_sequencing_of_FACS_sorted_cells_v2_/5829687. Cell assignments were acquired from the 'free_annotation' and 'subtissue' column of the metadata, to provide cells with either a cell identity or at the least a sub tissue identity. The core workflow was run once for all the cells and then again for cells from only the pancreas to allow direct comparisons to other datasets. Genes were filtered to 20 cells for each analysis.

Pancreas (Baron et al., 2016)

The Pancreas was followed up independently after found as an enriched tissue in adults through both datasets. Baron et al. (2016) sequenced 2000 pancreatic cells from 5 mice and identified 13 cell clusters including the main islets (alpha, beta, etc.). Raw data and cell metadata were acquired together for the mouse data from two files from Gene Expression Omnibus through accession no. GSE84133. The expression matrix included cell cluster labels through the 'assigned_cluster' annotations. Data were normalised to 10,000 UMI and log transformed and run through the core workflow once for the major cell types.

Muscle cells (De Micheli et al., 2020)

Muscle tissue was followed up independently after found as an enriched tissue in adult in the TM dataset. De Micheli et al., (2020) sequenced cells from injured tibalis anterior muscles at 0-, 2-, 5- or 7-days post injury (notexin injection). The non FACs sorted sample was used for this analysis and the raw, processed and metadata were acquired from Gene Expression Omnibus through accession no. GSE143437. Metadata included cell identities as 'cell-annotation' annotation. Raw data were used to apply the 20-cell filter to the genes and the processed data were filtered and run through the workflow once for all major cell types.

Mammary Gland Epithelial Cells (Bach et al., 2017)

The mammary gland, although not identified as a tissue of interest from the cross-tissue analysis, has been raised as a potential site of imprinted gene enrichment, particularly in the context of marsupials and very recently restated as a site of imprinting in mammals generally. Bach et al. (2017) use single-cell RNA sequencing to characterize the mammary epithelial cells across four developmental stages of the mammary gland (nulliparous, mid gestation, lactation, and post involution). Processed data and metadata were acquired through personal correspondence with the authors (although raw data is available from Gene Expression Omnibus through accession no. GSE106273). The matrix was scaled using the supplied size factors and log2 normalised. Cell annotations were taken from the 'SuperCluster' annotations involving 8 unique cell clusters and the 'Condition' annotations for developmental stage and cells were run through the workflow twice, once for cell types and once for developmental stage.

#### Supplemental Table S1

Custom List of Imprinted Genes used as the gene set in this study
('Supplemental_Table_S1_Imprinted_Gene_List.xlsx')

#### Supplemental Table S2

**Supplemental Table S2.** Imprinted gene over-representation in MCA adult cell populations (Han et al., 2018). *Identity* – Cell identities for the cells used in analysis; *Up Reg* – number of upregulated genes (total number of genes in the dataset in brackets number of imprinted genes upregulated with 2FC (total number of IGs in the dataset in brackets); *ORA p* – *p* value from over representation analysis on groups with minimum 5% of total IGs; *ORA q* – Bonferroni corrected *p* value from ORA; *Mean FC IG* – mean fold change for upregulated imprinted genes; *Mean FC Rest* – mean fold change for all other upregulated genes; *GSEA p* – *p* value from Gene Set Enrichment Analysis for identity groups with 15+ IGs and Mean FC IG > Mean FC Rest; *GSEA q* – Bonferroni corrected *p* values from GSEA.

| Tissue Cell Identity | Up Reg (20,534) | IG (95) | ORA *p* | ORA *q* | Mean FC IG | Mean FC Rest | GSEA *p* | GSEA q |
|---|---|---|---|---|---|---|---|---|
| **Stromal cell_Dpt high(Bladder)** | 1697 | 24 | 5.44E-07 | **4.90E-05** | 5.51 | 5.03 | 0.1514 | 0.6056 |
| **Endocrine cell(Pancreas)** | 666 | 14 | 2.25E-06 | **0.0002** | 12.06 | 15.68 | - | - |
| **Stromal cell_Smoc2 high(Pancreas)** | 1593 | 22 | 2.68E-06 | **0.0002** | 9.45 | 5.36 | 0.0673 | 0.2692 |
| **Stromal cell_Mfap4 high(Pancreas)** | 1164 | 18 | 5.57E-06 | **0.0005** | 6.08 | 7.56 | - | - |
| **Î²-cell(Pancreas)** | 1413 | 19 | 2.14E-05 | **0.0019** | 11.14 | 12.18 | - | - |
| **Stromal cell_Dcn high(Lung)** | 762 | 13 | 4.82E-05 | **0.0043** | 4.88 | 6.92 | - | - |
| **Stromal cell_Col3a1 high(Mammary-Gland-Virgin)** | 802 | 13 | 8.10E-05 | **0.0073** | 9.09 | 5.42 | - | - |
| **Acinar cell(Pancreas)** | 260 | 7 | 0.0002 | **0.0185** | 26.39 | 47.94 | - | - |
| **Stromal cell_Pi16 high(Mammary-Gland-Virgin)** | 655 | 11 | 0.0002 | **0.0198** | 6.83 | 8.03 | - | - |
| **Dividing cell(Pancreas)** | 1054 | 14 | 0.0003 | **0.0305** | 17.2 | 15.21 | - | - |
| **Mesenchymal stromal cell(Bladder)** | 1056 | 14 | 0.0003 | **0.0311** | 4.89 | 5.69 | - | - |
| **Myelinating oligodendrocyte(Brain)** | 2566 | 24 | 0.0005 | 0.0452 | 6.94 | 12.54 | - | - |
| **Ductal cell(Pancreas)** | 1878 | 19 | 0.0009 | 0.0782 | 5.24 | 6.07 | - | - |
| **Stromal cell_Car3 high(Bladder)** | 1316 | 15 | 0.001 | 0.091 | 6.66 | 6.56 | 0.3626 | 1 |
| Stromal cell_Has1 high(Uterus) | 2378 | 22 | 0.0011 | 0.0951 | 5.4 | 4.97 | 0.2102 | 0.8408 |
| Muscle progenitor cell(Muscle) | 257 | 6 | 0.0012 | 0.112 | 25.04 | 16.2 | - | - |
| **Stromal cell_Fn1 high(Pancreas)** | 1277 | 14 | 0.0022 | 0.1954 | 6.16 | 5.64 | - | - |
| **Stromal cell_Cxcl14 high(Uterus)** | 502 | 8 | 0.0023 | 0.2057 | 6.47 | 6.59 | - | - |
| **Stroma cell (Ovary)** | 1015 | 12 | 0.0025 | 0.2234 | 5.19 | 4.21 | - | - |
| **Oligodendrocyte precursor cell(Brain)** | 1756 | 17 | 0.0027 | 0.2423 | 9.36 | 16.09 | - | - |
| **Astroglial cell(Bergman glia)(Brain)** | 1113 | 12 | 0.0052 | 0.4681 | 6.71 | 21.88 | - | - |
| **Stromal cell_Ccl11 high(Uterus)** | 2750 | 22 | 0.0065 | 0.5855 | 4.37 | 4.87 | - | - |
| **Stromal cell_Hsd11b2 high(Uterus)** | 1235 | 12 | 0.0115 | 1 | 4 | 4.73 | - | - |
| **Neuron(Brain)** | 815 | 9 | 0.0131 | 1 | 33.07 | 30.25 | - | - |
| **Endothelial cell(Liver)** | 1128 | 11 | 0.015 | 1 | 3.69 | 5.42 | - | - |
| **Stromal cell_Dcn high(Kidney)** | 441 | 6 | 0.0166 | 1 | 10.65 | 13.16 | - | - |
| **Stromal cell(Muscle)** | 442 | 6 | 0.0168 | 1 | 12.65 | 15.26 | - | - |
| **Endothelial cell_Fabp4 high(Pancreas)** | 1148 | 11 | 0.0169 | 1 | 3.71 | 7.06 | - | - |
| **Schwann cell(Brain)** | 1017 | 10 | 0.0191 | 1 | 47.54 | 18.71 | - | - |
| **Erythroblast_Hbb-bt high(Pancreas)** | 460 | 6 | 0.02 | 1 | 7.8 | 18.78 | - | - |
| **Macrophage_Klf2 high(Brain)** | 1080 | 10 | 0.0276 | 1 | 5.51 | 7.85 | - | - |
| **Pan-GABAergic(Brain)** | 943 | 9 | 0.0304 | 1 | 11.17 | 17.93 | - | - |
| **Granulocyte(Uterus)** | 527 | 6 | 0.0356 | 1 | 5.41 | 9.04 | - | - |
| **Muscle cell_Mgp high(Uterus)** | 676 | 7 | 0.0373 | 1 | 4.69 | 5.67 | - | - |
| **Stromal cell_Inmt high(Lung)** | 677 | 7 | 0.0375 | 1 | 7.83 | 9.41 | - | - |
| **AT2 Cell(Lung)** | 1198 | 10 | 0.0501 | 1 | 3.92 | 7.68 | - | - |
| **Endothelial cell_Tm4sf1 high(Pancreas)** | 727 | 7 | 0.0516 | 1 | 7.86 | 8.82 | - | - |
| **S1 proximal tubule cells(Kidney)** | 753 | 7 | 0.06 | 1 | 8.32 | 11.72 | - | - |
| **Smooth muscle cell_Acta2 high(Pancreas)** | 460 | 5 | 0.0624 | 1 | 26.8 | 16.96 | - | - |
| **Stromal cell_Gm23935 high(Uterus)** | 761 | 7 | 0.0628 | 1 | 8.79 | 7.71 | - | - |
| **Proximal tubule cell_Osgin1 high(Kidney)** | 935 | 8 | 0.0678 | 1 | 11.61 | 10.13 | - | - |
| **Granulosa cell_Kctd14 high(Ovary)** | 631 | 6 | 0.0724 | 1 | 5.46 | 6.51 | - | - |
| **Cumulus cell_Ube2c high(Ovary)** | 2016 | 14 | 0.0801 | 1 | 4.4 | 4.31 | - | - |
| **Sertoli cell(Testis)** | 1327 | 10 | 0.0863 | 1 | 17.91 | 18.39 | - | - |
| **Astrocyte_Mfe8 high(Brain)** | 987 | 8 | 0.0865 | 1 | 5.67 | 15.78 | - | - |
| **AT1 Cell(Lung)** | 669 | 6 | 0.0899 | 1 | 5.55 | 11.15 | - | - |
| **Hypothalamic ependymal cell(Brain)** | 1005 | 8 | 0.0936 | 1 | 23.6 | 36.13 | - | - |
| **Granulosa cell_Inhba high(Ovary)** | 684 | 6 | 0.0974 | 1 | 4.45 | 6.46 | - | - |
| **Ovarian surface epithelium cell(Ovary)** | 1017 | 8 | 0.0986 | 1 | 11.42 | 10.08 | - | - |
| **Epithelial cell_Gkn3 high(Stomach)** | 943 | 7 | 0.1465 | 1 | 8.55 | 9.04 | - | - |

| | | | | | | | | |
|---|---|---|---|---|---|---|---|---|
| **Basal epithelial cell(Bladder)** | 2621 | 16 | 0.1494 | 1 | 4.05 | 4.79 | - | - |
| **Urothelium(Bladder)** | 4633 | 26 | 0.1584 | 1 | 4.18 | 5.03 | - | - |
| **Epithelial cell_Gm23935 high(Bladder)** | 789 | 6 | 0.1587 | 1 | 10.12 | 10.14 | - | - |
| **Ciliated cell(Lung)** | 1031 | 7 | 0.2001 | 1 | 11.17 | 19.91 | - | - |
| **Ovarian vascular surface endothelium cell(Ovary)** | 1852 | 11 | 0.236 | 1 | 4.48 | 4.76 | - | - |
| **Thecal cell(Ovary)** | 907 | 6 | 0.2429 | 1 | 3.65 | 6.1 | - | - |
| **Alveolar macrophage_Ear2 high(Lung)** | 732 | 5 | 0.2509 | 1 | 4.66 | 7.63 | - | - |
| **Smooth muscle cell(Bladder)** | 736 | 5 | 0.2544 | 1 | 5.7 | 11.86 | - | - |
| **Astrocyte_Atp1b2 high(Brain)** | 778 | 5 | 0.2917 | 1 | 10.98 | 19.72 | - | - |
| **Macrophage_Lyz2 high(Brain)** | 784 | 5 | 0.2971 | 1 | 6.34 | 10.6 | - | - |
| **Proximal tubule brush border cell(Kidney)** | 979 | 6 | 0.2998 | 1 | 5.46 | 8.53 | - | - |
| **Glandular epithelium_Ltf high(Uterus)** | 802 | 5 | 0.3135 | 1 | 7.11 | 9.13 | - | - |
| **S cell_Chgb high(Small-Intestine)** | 976 | 5 | 0.4734 | 1 | 22.73 | 23.74 | - | - |
| **Pit cell_Ifrd1 high(Stomach)** | 1401 | 7 | 0.4734 | 1 | 3.85 | 5 | - | - |
| **Spermatocyte_Slc2a3 high(Testis)** | 4009 | 19 | 0.4944 | 1 | 11.81 | 15.38 | - | - |
| **Epithelial cell_Krt20 high(Stomach)** | 1446 | 7 | 0.5073 | 1 | 4.79 | 5.2 | - | - |
| **Stomach cell_Muc5ac high(Stomach)** | 1149 | 5 | 0.6198 | 1 | 5.17 | 5.2 | - | - |
| **Antral mucous cell (Stomach)** | 1221 | 5 | 0.6733 | 1 | 4.09 | 5.42 | - | - |
| **Epithelium of small intestinal villi_Fabp6 high(Small-Intestine)** | 1493 | 6 | 0.6973 | 1 | 4.62 | 10.56 | - | - |
| **Stomach cell_Mt2 high(Stomach)** | 1505 | 6 | 0.7048 | 1 | 3.82 | 4.5 | - | - |
| **Pre-Sertoli cell_Ctsl high(Testis)** | 1302 | 5 | 0.7275 | 1 | 23.91 | 30.85 | - | - |
| **Preleptotene spermatogonia(Testis)** | 1787 | 7 | 0.7306 | 1 | 19.13 | 13.69 | - | - |
| **Large luteal cell(Ovary)** | 1585 | 6 | 0.7512 | 1 | 2.94 | 4.63 | - | - |
| **Spermatids_1700016P04Rik high(Testis)** | 1430 | 5 | 0.7997 | 1 | 5.59 | 13.52 | - | - |
| **Parietal cell (Stomach)** | 1928 | 7 | 0.7998 | 1 | 8.87 | 6.32 | - | - |
| **Epithelial cell_Kcne3 high(Small-Intestine)** | 1795 | 6 | 0.8478 | 1 | 18.45 | 4.72 | - | - |
| **Umbrella cell(Bladder)** | 6556 | 26 | 0.8572 | 1 | 4.65 | 5.76 | - | - |
| **Spermatocyte_Cabs1 high(Testis)** | 1690 | 5 | 0.9001 | 1 | 5.93 | 16.23 | - | - |
| **Spermatogonia_Tbc1d23 high(Testis)** | 3374 | 11 | 0.9274 | 1 | 15.93 | 28.02 | - | - |
| **Monocyte_Elane high(Peripheral_Blood)** | 2388 | 7 | 0.9357 | 1 | 4.73 | 4.45 | - | - |
| **luteal cells(Ovary)** | 4236 | 14 | 0.9443 | 1 | 8.63 | 4.15 | - | - |
| **Elongating spermatid(Testis)** | 2600 | 7 | 0.9647 | 1 | 4.69 | 10.87 | - | - |
| **Spermatocyte_Calm2 high(Testis)** | 2460 | 6 | 0.9772 | 1 | 14.99 | 11.59 | - | - |
| **Erythroblast(Spleen)** | 2766 | 7 | 0.9786 | 1 | 15.81 | 11.14 | - | - |
| **Spermatids_Cst13 high(Testis)** | 2806 | 7 | 0.9811 | 1 | 8.99 | 21.46 | - | - |
| **Spermatogonia_1700001P01Rik high(Testis)** | 2519 | 6 | 0.9811 | 1 | 5.46 | 17.48 | - | - |
| **Erythroblast_Car2 high(Peripheral_Blood)** | 2834 | 7 | 0.9826 | 1 | 4.19 | 5.83 | - | - |
| **Spermatocyte_1700001F09Rik high(Testis)** | 4544 | 13 | 0.9865 | 1 | 9.62 | 18.76 | - | - |
| **Erythroblast_Car1 high(Muscle)** | 2646 | 5 | 0.9958 | 1 | 10.82 | 6.32 | - | - |
| **Glandular epithelium_Sprr2f high(Uterus)** | 3352 | 7 | 0.9969 | 1 | 2.99 | 5.99 | - | - |
| **abT cell(Thymus)** | 648 | 2 | - | - | 2.98 | 5.55 | - | - |
| **Alveolar bipotent progenitor(Lung)** | 366 | 2 | - | - | 8.57 | 15.64 | - | - |
| **Alveolar macrophage_Pclaf high(Lung)** | 528 | 0 | - | - | 0 | 14.36 | - | - |
| **B Cell(Lung)** | 436 | 0 | - | - | 0 | 9.15 | - | - |
| **B cell(Thymus)** | 1360 | 4 | - | - | 4 | 4.05 | - | - |
| **B cell(Uterus)** | 525 | 2 | - | - | 6.77 | 16.17 | - | - |
| **B cell_Cd79a&Fcer2a high(Mammary-Gland-Virgin)** | 468 | 2 | - | - | 7.04 | 7.17 | - | - |
| **B cell_Cd79a&Iglc2 high(Mammary-Gland-Virgin)** | 458 | 1 | - | - | 3.06 | 5.43 | - | - |
| **B cell_Fcmr high(Liver)** | 568 | 2 | - | - | 4.63 | 8.85 | - | - |
| **B cell_Igha high(Peripheral_Blood)** | 480 | 4 | - | - | 4.84 | 13.03 | - | - |
| **B cell_Ighd high(Small-Intestine)** | 397 | 0 | - | - | 0 | 11.5 | - | - |
| **B cell_Igkc high(Bone-Marrow)** | 411 | 0 | - | - | 0 | 27.5 | - | - |
| **B cell_Igkv12-46 high(Small-Intestine)** | 487 | 2 | - | - | 4.51 | 14.95 | - | - |
| **B cell_Jchain high(Liver)** | 447 | 1 | - | - | 3.35 | 11.91 | - | - |
| **B cell_Jchain high(Muscle)** | 293 | 1 | - | - | 7.1 | 16.1 | - | - |
| **B cell_Jchain high(Small-Intestine)** | 416 | 4 | - | - | 4.14 | 12.82 | - | - |
| **B cell_Ly6d high(Peripheral_Blood)** | 540 | 3 | - | - | 6.36 | 4.74 | - | - |
| **B cell_Ms4a1 high(Small-Intestine)** | 1117 | 2 | - | - | 5.54 | 6.26 | - | - |
| **B cell_Rps27rt high(Peripheral_Blood)** | 388 | 0 | - | - | 0 | 17.52 | - | - |
| **B cell_Vpreb3 high(Muscle)** | 722 | 0 | - | - | 0 | 6.42 | - | - |
| **B cell_Vpreb3 high(Peripheral_Blood)** | 1076 | 2 | - | - | 3.08 | 5.1 | - | - |
| **Basophil_Prss34 high(Peripheral_Blood)** | 464 | 1 | - | - | 9.28 | 16.72 | - | - |
| **Clara Cell(Lung)** | 521 | 3 | - | - | 6.9 | 11.13 | - | - |
| **Columnar epithelium(Small-Intestine)** | 1523 | 3 | - | - | 4.13 | 8.39 | - | - |
| **Conventional dendritic cell_Gngt2 high(Lung)** | 404 | 1 | - | - | 24.93 | 10.64 | - | - |
| **Conventional dendritic cell_H2-M2 high(Lung)** | 283 | 0 | - | - | 0 | 22.95 | - | - |

| | | | | | | | | |
|---|---|---|---|---|---|---|---|---|
| **Conventional dendritic cell_Mgl2 high(Lung)** | 418 | 1 | - | - | 6.19 | 10.63 | - | - |
| **Conventional dendritic cell_Tubb5 high(Lung)** | 434 | 0 | - | - | 0 | 12.87 | - | - |
| **Cumulus cell_Car14 high(Ovary)** | 1321 | 4 | - | - | 3.5 | 4.34 | - | - |
| **Cumulus cell_Nupr1 high(Ovary)** | 527 | 4 | - | - | 6.11 | 5.55 | - | - |
| **Dendritic cell(Prostate)** | 102 | 0 | - | - | 0 | 15.91 | - | - |
| **Dendritic cell(Stomach)** | 670 | 3 | - | - | 7.01 | 10.39 | - | - |
| **Dendritic cell(Uterus)** | 459 | 0 | - | - | 0 | 8.25 | - | - |
| **Dendritic cell_Ccr7 high(Kidney)** | 227 | 0 | - | - | 0 | 15.56 | - | - |
| **Dendritic cell_Cd74 high(Bladder)** | 403 | 0 | - | - | 0 | 9.7 | - | - |
| **Dendritic cell_Cst3 high(Kidney)** | 207 | 0 | - | - | 0 | 16.37 | - | - |
| **Dendritic cell_Cst3 high(Liver)** | 820 | 2 | - | - | 3.5 | 5.26 | - | - |
| **Dendritic cell_Cst3 high(Mammary-Gland-Virgin)** | 457 | 1 | - | - | 6.33 | 7.96 | - | - |
| **Dendritic cell_Fscn1 high(Mammary-Gland-Virgin)** | 723 | 2 | - | - | 7.43 | 8.04 | - | - |
| **Dendritic cell_Lyz2 high(Bladder)** | 643 | 1 | - | - | 7.19 | 8.98 | - | - |
| **Dendritic cell_Naaa high(Lung)** | 438 | 0 | - | - | 0 | 9.92 | - | - |
| **Dendritic cell_S100a4 high(Spleen)** | 597 | 2 | - | - | 4.25 | 8.83 | - | - |
| **Dendritic cell_Siglech high(Liver)** | 598 | 2 | - | - | 8.07 | 8.95 | - | - |
| **Dendritic cell_Siglech high(Mammary-Gland-Virgin)** | 319 | 1 | - | - | 2.53 | 15.78 | - | - |
| **Dendritic cell_Siglech high(Peripheral_Blood)** | 502 | 1 | - | - | 38.3 | 8.7 | - | - |
| **Dendritic cell_Siglech high(Small-Intestine)** | 522 | 3 | - | - | 8.5 | 14.82 | - | - |
| **Dendritic cell_Siglech high(Spleen)** | 431 | 2 | - | - | 9.46 | 17.39 | - | - |
| **Dendrtic cell(Pancreas)** | 398 | 0 | - | - | 0 | 12.44 | - | - |
| **Dendrtic cell_Cst3 high(Small-Intestine)** | 390 | 0 | - | - | 0 | 10.42 | - | - |
| **Distal collecting duct principal cell_Cldn4 high(Kidney)** | 471 | 3 | - | - | 7.84 | 12.46 | - | - |
| **Distal collecting duct principal cell_Hsd11b2 high(Kidney)** | 453 | 3 | - | - | 19.82 | 13.29 | - | - |
| **Distal convoluted tubule_Pvalb high(Kidney)** | 642 | 2 | - | - | 4.61 | 8.54 | - | - |
| **Distal convoluted tubule_S100g high(Kidney)** | 453 | 1 | - | - | 6.59 | 11.55 | - | - |
| **Dividing cell(Mammary-Gland-Virgin)** | 1434 | 1 | - | - | 3.37 | 4.09 | - | - |
| **Dividing cell(Stomach)** | 2065 | 4 | - | - | 3.3 | 4.2 | - | - |
| **Dividing cells(Lung)** | 683 | 4 | - | - | 6.68 | 11.83 | - | - |
| **Dividing dendritic cells(Lung)** | 509 | 1 | - | - | 3.58 | 13.66 | - | - |
| **Dividing T cells(Lung)** | 668 | 0 | - | - | 0 | 8.52 | - | - |
| **DPT cell(Thymus)** | 1061 | 3 | - | - | 3.96 | 5.05 | - | - |
| **Endothelial cell(Kidney)** | 632 | 1 | - | - | 4.46 | 13.82 | - | - |
| **Endothelial cell(Muscle)** | 215 | 1 | - | - | 2.4 | 19.08 | - | - |
| **Endothelial cell_Cldn5 high(Uterus)** | 424 | 4 | - | - | 16.15 | 27.76 | - | - |
| **Endothelial cell_Kdr high(Lung)** | 436 | 3 | - | - | 13.93 | 16.66 | - | - |
| **Endothelial cell_Lrg1 high(Pancreas)** | 521 | 4 | - | - | 6.02 | 15.13 | - | - |
| **Endothelial cell_Ly6c1 high(Bladder)** | 691 | 3 | - | - | 28.86 | 11.41 | - | - |
| **Endothelial cell_Tm4sf1 high(Uterus)** | 701 | 4 | - | - | 4.11 | 6.29 | - | - |
| **Endothelial cell_Tmem100 high(Lung)** | 491 | 1 | - | - | 5.95 | 13.81 | - | - |
| **Endothelial cells_Vwf high(Lung)** | 506 | 4 | - | - | 26.6 | 15.42 | - | - |
| **Epithelia cell_Spp1 high(Liver)** | 354 | 1 | - | - | 5.52 | 15.66 | - | - |
| **Epithelial cell(Liver)** | 464 | 3 | - | - | 14.4 | 21.76 | - | - |
| **Epithelial cell(Prostate)** | 158 | 0 | - | - | 0 | 7.43 | - | - |
| **Epithelial cell_Cryab high(Kidney)** | 460 | 3 | - | - | 6.39 | 14.63 | - | - |
| **Epithelial cell_Sh2d6 high(Small-Intestine)** | 715 | 1 | - | - | 8.15 | 16.77 | - | - |
| **Epithelial cell_Upk3a high(Bladder)** | 766 | 4 | - | - | 4.33 | 7.46 | - | - |
| **Epithelium of small intestinal villi_Fabp1 high(Small-Intestine)** | 1488 | 3 | - | - | 5.47 | 9.1 | - | - |
| **Epithelium of small intestinal villi_Gm23935 high(Small-Intestine)** | 585 | 4 | - | - | 12.89 | 20.42 | - | - |
| **Epithelium of small intestinal villi_S100g high(Small-Intestine)** | 469 | 2 | - | - | 7.2 | 26.76 | - | - |
| **Erythroblast_Car2 high(Muscle)** | 677 | 3 | - | - | 4.07 | 15.44 | - | - |
| **Erythroblast_Hbb-bs high(Liver)** | 721 | 3 | - | - | 4.51 | 6.22 | - | - |
| **Erythroblast_Hbb-bs high(Testis)** | 690 | 2 | - | - | 38.58 | 35.53 | - | - |
| **Erythroblast_Hbb-bt high(Liver)** | 269 | 2 | - | - | 6.39 | 22.25 | - | - |
| **Erythroblast_Igkc high(Pancreas)** | 289 | 4 | - | - | 5.11 | 21.85 | - | - |
| **Fenestrated endothelial cell_Plvap high(Kidney)** | 433 | 4 | - | - | 6.78 | 12.15 | - | - |

| | | | | | | | | |
|---|---|---|---|---|---|---|---|---|
| **Fenestrated endothelial cell_Tm4sf1 high(Kidney)** | 264 | 1 | - | - | 2.3 | 17.88 | - | - |
| **G cell(Stomach)** | 538 | 2 | - | - | 6.11 | 19.02 | - | - |
| **Gastric mucosal cell(Stomach)** | 743 | 3 | - | - | 4.65 | 6.95 | - | - |
| **gdT cell (Thymus)** | 752 | 1 | - | - | 34.92 | 7.79 | - | - |
| **Glandular epithelium(Prostate)** | 182 | 1 | - | - | 3.35 | 9.83 | - | - |
| **Granulocyte monocyte progenitor cell(Muscle)** | 1053 | 4 | - | - | 5.95 | 5.98 | - | - |
| **Granulocyte(Liver)** | 707 | 3 | - | - | 14.51 | 8.57 | - | - |
| **Granulocyte(Spleen)** | 303 | 0 | - | - | 0 | 16.32 | - | - |
| **Hematopoietic stem progenitor cell(Bone-Marrow)** | 998 | 3 | - | - | 6.54 | 21.02 | - | - |
| **Hepatocyte_Fabp1 high(Liver)** | 519 | 1 | - | - | 4.25 | 28.72 | - | - |
| **Intercalated cells of collecting duct_Aqp6 high(Kidney)** | 580 | 1 | - | - | 4.3 | 11.87 | - | - |
| **Intercalated cells of collecting duct_Slc26a4 high(Kidney)** | 567 | 3 | - | - | 4.33 | 8.79 | - | - |
| **Interstitial macrophage(Lung)** | 520 | 1 | - | - | 7.06 | 7.78 | - | - |
| **Keratinocyte(Uterus)** | 407 | 4 | - | - | 9.58 | 19.75 | - | - |
| **Kuppfer cell(Liver)** | 1274 | 4 | - | - | 3.36 | 5.59 | - | - |
| **Leydig cell(Testis)** | 1533 | 4 | - | - | 5.6 | 15.4 | - | - |
| **Luminal cell_Krt19 high (Mammary-Gland-Virgin)** | 1046 | 4 | - | - | 3.51 | 6.47 | - | - |
| **Luminal progenitor(Mammary-Gland-Virgin)** | 606 | 2 | - | - | 16.45 | 10.88 | - | - |
| **Macrophage(Pancreas)** | 925 | 3 | - | - | 4.32 | 5.34 | - | - |
| **Macrophage(Spleen)** | 556 | 2 | - | - | 5.25 | 14.09 | - | - |
| **Macrophage(Stomach)** | 219 | 0 | - | - | 0 | 20.14 | - | - |
| **Macrophage(Uterus)** | 887 | 2 | - | - | 2.42 | 5.02 | - | - |
| **Macrophage_Ace high(Peripheral_Blood)** | 522 | 1 | - | - | 3.39 | 6.38 | - | - |
| **Macrophage_Apoe high(Small-Intestine)** | 420 | 1 | - | - | 2.02 | 13.64 | - | - |
| **Macrophage_C1qc high(Mammary-Gland-Virgin)** | 589 | 1 | - | - | 3.82 | 8.46 | - | - |
| **Macrophage_Ccl4 high (Kidney)** | 402 | 1 | - | - | 3.93 | 11.23 | - | - |
| **Macrophage_Chil3 high(Liver)** | 689 | 1 | - | - | 5.69 | 5.94 | - | - |
| **Macrophage_Flt-ps1 high(Peripheral_Blood)** | 659 | 1 | - | - | 28.77 | 11.89 | - | - |
| **Macrophage_G0s2 high(Small-Intestine)** | 414 | 2 | - | - | 28.11 | 17.9 | - | - |
| **Macrophage_Ly6c2 high(Pancreas)** | 352 | 1 | - | - | 8.69 | 17.62 | - | - |
| **Macrophage_Lyz1 high(Mammary-Gland-Virgin)** | 594 | 0 | - | - | 0 | 5 | - | - |
| **Macrophage_Lyz2 high(Ovary)** | 807 | 1 | - | - | 6.58 | 6.08 | - | - |
| **Macrophage_Lyz2 high(Testis)** | 1271 | 4 | - | - | 11.52 | 22.85 | - | - |
| **Macrophage_Ms4a6c high(Bone-Marrow)** | 1074 | 1 | - | - | 5.97 | 21.19 | - | - |
| **Macrophage_Ms4a6c high(Muscle)** | 846 | 2 | - | - | 4.82 | 5.22 | - | - |
| **Macrophage_Pf4 high(Bladder)** | 612 | 1 | - | - | 9.79 | 11.71 | - | - |
| **Macrophage_Retnla high(Muscle)** | 445 | 0 | - | - | 0 | 11.6 | - | - |
| **Macrophage_S100a4 high(Peripheral_Blood)** | 802 | 2 | - | - | 3.02 | 4.42 | - | - |
| **Marginal zone B cell(Spleen)** | 1009 | 1 | - | - | 5.67 | 5.38 | - | - |
| **Microglia(Brain)** | 644 | 3 | - | - | 4.25 | 8.13 | - | - |
| **Monocyte progenitor cell(Lung)** | 604 | 0 | - | - | 0 | 14.88 | - | - |
| **Monocyte(Spleen)** | 642 | 1 | - | - | 4 | 7.53 | - | - |
| **Monocyte(Uterus)** | 543 | 3 | - | - | 6.69 | 7.77 | - | - |
| **Monocyte_F13a1 high(Peripheral_Blood)** | 1085 | 3 | - | - | 3.16 | 4.81 | - | - |
| **Muscle cell(Stomach)** | 424 | 3 | - | - | 6.39 | 14.03 | - | - |
| **Muscle cell_Pcp4 high(Uterus)** | 302 | 3 | - | - | 21.03 | 22.24 | - | - |
| **Muscle cell_Tnnc2 high(Muscle)** | 350 | 3 | - | - | 88.46 | 63.03 | - | - |
| **Neutrophil granulocyte(Lung)** | 309 | 0 | - | - | 0 | 20.84 | - | - |
| **Neutrophil progenitor(Bone-Marrow)** | 409 | 2 | - | - | 17.38 | 20.3 | - | - |
| **Neutrophil(Spleen)** | 534 | 0 | - | - | 0 | 16.81 | - | - |
| **Neutrophil_Camp high(Muscle)** | 546 | 1 | - | - | 3.88 | 11.44 | - | - |
| **Neutrophil_Camp high(Peripheral_Blood)** | 979 | 4 | - | - | 3.59 | 6.06 | - | - |
| **Neutrophil_Il1b high(Peripheral_Blood)** | 510 | 4 | - | - | 5.48 | 9.86 | - | - |
| **Neutrophil_Ltf high(Peripheral_Blood)** | 455 | 2 | - | - | 21.34 | 12.88 | - | - |
| **Neutrophil_Mmp8 high(Bone-Marrow)** | 514 | 1 | - | - | 22.67 | 20.6 | - | - |
| **Neutrophil_Ngp high(Bone-Marrow)** | 513 | 3 | - | - | 6.79 | 21.02 | - | - |
| **Neutrophil_Ngp high(Liver)** | 274 | 0 | - | - | 0 | 24.2 | - | - |
| **Neutrophil_Retnlg high(Peripheral_Blood)** | 304 | 1 | - | - | 212.78 | 21.4 | - | - |
| **NK cell(Bladder)** | 468 | 1 | - | - | 8.45 | 13.71 | - | - |
| **NK Cell(Lung)** | 343 | 0 | - | - | 0 | 14.5 | - | - |
| **NK cell(Mammary-Gland-Virgin)** | 380 | 0 | - | - | 0 | 11.4 | - | - |

| | | | | | | | | |
|---|---|---|---|---|---|---|---|---|
| **NK cell(Spleen)** | 379 | 1 | - | - | 6.34 | 11.19 | - | - |
| **NK cell(Uterus)** | 951 | 3 | - | - | 10 | 6.5 | - | - |
| **NK cell_Gzma high(Peripheral_Blood)** | 446 | 2 | - | - | 5.08 | 8.06 | - | - |
| **Nuocyte(Lung)** | 277 | 1 | - | - | 82.97 | 16.1 | - | - |
| **Paneth cell(Small-Intestine)** | 716 | 4 | - | - | 13.23 | 27.1 | - | - |
| **Pericentral (PC) hepatocytes(Liver)** | 652 | 3 | - | - | 14.43 | 25.01 | - | - |
| **Periportal (PP) hepatocyte(Liver)** | 705 | 4 | - | - | 29.62 | 57.64 | - | - |
| **Pit cell_Gm26917 high(Stomach)** | 635 | 3 | - | - | 7.29 | 14.34 | - | - |
| **Plasma cell(Spleen)** | 655 | 2 | - | - | 4.2 | 11.26 | - | - |
| **Plasmacytoid dendritic cell(Lung)** | 479 | 1 | - | - | 22.07 | 5.5 | - | - |
| **Pre-pro B cell(Bone-Marrow)** | 863 | 0 | - | - | 0 | 14.94 | - | - |
| **Pre-Sertoli cell_Cst9 high(Testis)** | 1621 | 3 | - | - | 10.66 | 14.58 | - | - |
| **Proliferating thymocyte(Thymus)** | 1614 | 4 | - | - | 2.87 | 4.68 | - | - |
| **Prostate gland cell(Prostate)** | 270 | 0 | - | - | 0 | 12.43 | - | - |
| **Proximal tubule cell_Cyp4a14 high(Kidney)** | 610 | 4 | - | - | 9.41 | 14.54 | - | - |
| **S cell_Gip high(Small-Intestine)** | 413 | 3 | - | - | 25.8 | 26.24 | - | - |
| **S3 proximal tubule cells(Kidney)** | 989 | 3 | - | - | 24.91 | 14.17 | - | - |
| **Small luteal cell(Ovary)** | 385 | 1 | - | - | 27.16 | 5.05 | - | - |
| **Smooth muscle cell_Rgs5 high(Pancreas)** | 342 | 3 | - | - | 7.22 | 23.46 | - | - |
| **Smooth muscle cell_Rgs5 high(Uterus)** | 257 | 2 | - | - | 18.96 | 22.72 | - | - |
| **Spermatids_Hmgb4 high(Testis)** | 1949 | 2 | - | - | 42.32 | 24.78 | - | - |
| **Spermatids_Tnp1 high(Testis)** | 2092 | 4 | - | - | 9.11 | 16.5 | - | - |
| **Spermatocyte_Mesp1 high(Testis)** | 2113 | 3 | - | - | 6.74 | 11.42 | - | - |
| **Stem and progenitor cell(Mammary-Gland-Virgin)** | 466 | 0 | - | - | 0 | 8.52 | - | - |
| **Stomach cell_Gkn2 high(Stomach)** | 765 | 2 | - | - | 3.86 | 4.79 | - | - |
| **Stromal cell(Liver)** | 375 | 3 | - | - | 15.53 | 23.18 | - | - |
| **Stromal cell(Prostate)** | 129 | 0 | - | - | 0 | 12.11 | - | - |
| **Stromal cell_Acta2 high(Lung)** | 326 | 2 | - | - | 4.08 | 18.59 | - | - |
| **Stromal cell_Adamdec1 high(Small-Intestine)** | 611 | 4 | - | - | 8.21 | 17.09 | - | - |
| **Stromal cell_Ankrd1 high(Kidney)** | 265 | 2 | - | - | 36.98 | 20.72 | - | - |
| **Stromal cell_Cxcl10 high(Kidney)** | 349 | 0 | - | - | 0 | 18 | - | - |
| **Stromal cell_Dcn high(Small-Intestine)** | 682 | 4 | - | - | 13.96 | 17.59 | - | - |
| **Stromal cell_Mgp high(Kidney)** | 274 | 0 | - | - | 0 | 17.54 | - | - |
| **Stromal cell_Ptgds high(Kidney)** | 423 | 3 | - | - | 10.59 | 11.37 | - | - |
| **T-cells_Ctla4 high(Mammary-Gland-Virgin)** | 542 | 1 | - | - | 6.08 | 6.37 | - | - |
| **T cell(Kidney)** | 314 | 4 | - | - | 14.33 | 13.81 | - | - |
| **T cell(Pancreas)** | 196 | 2 | - | - | 28.57 | 24.42 | - | - |
| **T cell(Prostate)** | 128 | 0 | - | - | 0 | 12.81 | - | - |
| **T cell(Spleen)** | 907 | 1 | - | - | 5.5 | 4.2 | - | - |
| **T cell_Ccl5 high(Small-Intestine)** | 555 | 0 | - | - | 0 | 6.62 | - | - |
| **T cell_Cd7 high(Small-Intestine)** | 1004 | 4 | - | - | 4.29 | 6.5 | - | - |
| **T Cell_Cd8b1 high(Lung)** | 293 | 0 | - | - | 0 | 12.16 | - | - |
| **T cell_Cd8b1 high(Mammary-Gland-Virgin)** | 628 | 1 | - | - | 3.41 | 4.15 | - | - |
| **T cell_Gm14303 high(Peripheral_Blood)** | 648 | 1 | - | - | 3.85 | 10.7 | - | - |
| **T cell_Gzma high(Liver)** | 531 | 3 | - | - | 24.33 | 12.02 | - | - |
| **T cell_Icos high(Small-Intestine)** | 838 | 2 | - | - | 4.06 | 6.29 | - | - |
| **T cell_Id2 high(Thymus)** | 594 | 2 | - | - | 5.06 | 12.02 | - | - |
| **T cell_Ly6c2 high(Mammary-Gland-Virgin)** | 527 | 0 | - | - | 0 | 4.94 | - | - |
| **T cell_Ms4a4b high(Bone-Marrow)** | 518 | 2 | - | - | 16.65 | 19.15 | - | - |
| **T cell_Ms4a4b high(Small-Intestine)** | 494 | 1 | - | - | 24.05 | 13.92 | - | - |
| **T cell_Ms4a4b high(Thymus)** | 907 | 3 | - | - | 5.28 | 4.46 | - | - |
| **T cell_Trbc2 high(Liver)** | 715 | 4 | - | - | 5.4 | 6.28 | - | - |
| **T cell_Trbc2 high(Peripheral_Blood)** | 511 | 1 | - | - | 4.66 | 4.32 | - | - |
| **Thick ascending limb of the loop of Henle(Kidney)** | 605 | 1 | - | - | 3.8 | 13.46 | - | - |
| **Tuft cell(Stomach)** | 983 | 4 | - | - | 8.23 | 10.84 | - | - |
| **Ureteric epithelium(Kidney)** | 384 | 0 | - | - | 0 | 13.68 | - | - |
| **Vascular endothelial cell(Bladder)** | 742 | 1 | - | - | 8.31 | 8.35 | - | - |
| **Vascular smooth muscle progenitor cell(Bladder)** | 351 | 3 | - | - | 19.37 | 26.2 | - | - |

### Supplemental Table S3

**Supplemental Table S3.** Imprinted gene over-representation in Tabula Muris adult cell populations (Schaum et al., 2018). Identity – Cell identities for the cells used in analysis; All other column descriptions can be found in the legend of Table S2.

| Tissue Cell Identity | Up Reg (20,839) | IG (107) | ORA *p* | ORA *q* | Mean FC IG | Mean FC Rest | GSEA *p* | GSEA q |
|---|---|---|---|---|---|---|---|---|
| **skeletal muscle satellite stem cell (Diaphragm)** | 462 | 24 | 1.01E-17 | **5.65E-16** | 9.22 | 6.94 | 0.3174 | 1 |
| **skeletal muscle satellite cell (Limb Muscle)** | 833 | 23 | 3.05E-11 | **1.71E-09** | 16.53 | 6.91 | 0.0553 | 0.7742 |
| **mesenchymal stem cell (Diaphragm & Limb Muscle)** | 1289 | 24 | 2.78E-08 | **1.56E-06** | 4.69 | 4.25 | 0.1576 | 1 |
| **pancreatic stellate cell** | 1953 | 28 | 3.89E-07 | **2.18E-05** | 4.75 | 6.03 | - | - |
| **mesenchymal cell (Trachae)** | 2500 | 32 | 5.80E-07 | **3.25E-05** | 4.71 | 4.91 | - | - |
| **stromal cell (Mammary Gland & Lung)** | 1528 | 24 | 6.48E-07 | **3.63E-05** | 3.43 | 4.2 | - | - |
| **mesenchymal stem cell of adipose** | 2396 | 30 | 2.35E-06 | **0.0001** | 5.04 | 4.77 | 0.4084 | 1 |
| **pancreatic D cell** | 4713 | 45 | 5.54E-06 | **0.0003** | 11.57 | 7.89 | 0.0932 | 1 |
| **mesenchymal cell (Bladder)** | 2820 | 30 | 5.86E-05 | **0.0033** | 4.62 | 5.79 | - | - |
| **neuron (Brain Non-Myeloid)** | 5492 | 47 | 6.22E-05 | **0.0035** | 15.58 | 16.95 | - | - |
| **endocardial cell (Heart)** | 912 | 15 | 6.46E-05 | **0.0036** | 11.37 | 5.68 | 0.0146 | 0.2044 |
| **type B pancreatic cell** | 5024 | 44 | 7.28E-05 | **0.0041** | 14.61 | 8.89 | 0.0151 | 0.2114 |
| **oligodendrocyte precursor cell (Brain Non-Myeloid)** | 3969 | 36 | 0.0002 | **0.0134** | 6.69 | 12.41 | - | - |
| **pancreatic PP cell** | 4385 | 38 | 0.0004 | **0.0215** | 8.64 | 6.64 | 0.1114 | 1 |
| **oligodendrocyte (Brain Non-Myeloid)** | 1860 | 21 | 0.0004 | **0.0252** | 7.37 | 11.7 | - | - |
| **pancreatic A cell** | 4179 | 35 | 0.0014 | 0.0776 | 11.43 | 7.17 | 0.0069 | 0.0966 |
| smooth muscle cell | 968 | 12 | 0.004 | 0.2266 | 3.31 | 10.68 | - | - |
| **cardiac neuron** | 874 | 11 | 0.0053 | 0.2951 | 4.71 | 12.44 | - | - |
| **myofibroblast cell** | 1010 | 11 | 0.0147 | 0.8205 | 3.81 | 7.45 | - | - |
| **astrocyte of the cerebral cortex** | 1958 | 10 | 0.55559 | 1 | 4.67 | 12.44 | - | - |
| **basal cell** | 2187 | 10 | 0.697375 | 1 | 4.38 | 5.59 | - | - |
| **basal cell of epidermis** | 3100 | 11 | 0.936016 | 1 | 4.37 | 4.91 | - | - |
| **basophil** | 2286 | 7 | 0.956529 | 1 | 5.24 | 9.55 | - | - |
| **Bergmann glial cell** | 1848 | 15 | 0.050516 | 1 | 3.46 | 11.93 | - | - |
| **bladder urothelial cell** | 3858 | 17 | 0.793197 | 1 | 4.05 | 5.98 | - | - |
| **brain pericyte** | 1761 | 14 | 0.066599 | 1 | 3.76 | 8.94 | - | - |
| **Brush cell of epithelium proper of large intestine** | 2232 | 7 | 0.948583 | 1 | 5.43 | 23.92 | - | - |
| **cardiac muscle cell** | 2801 | 12 | 0.790389 | 1 | 4.01 | 19.7 | - | - |
| **ciliated columnar cell of tracheobronchial tree** | 2757 | 16 | 0.339523 | 1 | 10.07 | 31.52 | - | - |
| **common lymphoid progenitor** | 5746 | 24 | 0.905771 | 1 | 4.86 | 4.32 | 0.5291 | 1 |
| **endothelial cell** | 1017 | 8 | 0.151911 | 1 | 3.22 | 7.25 | - | - |
| **endothelial cell of hepatic sinusoid** | 1144 | 9 | 0.133479 | 1 | 5.48 | 9.63 | - | - |
| **enteroendocrine cell** | 2399 | 11 | 0.69994 | 1 | 13.32 | 12.81 | - | - |
| **epidermal cell** | 2335 | 9 | 0.860241 | 1 | 4.35 | 7.02 | - | - |
| **epithelial cell** | 1348 | 9 | 0.255566 | 1 | 5.49 | 10.57 | - | - |
| **epithelial cell of large intestine** | 3733 | 13 | 0.959638 | 1 | 5.77 | 5.17 | - | - |
| **epithelial cell of lung** | 1389 | 13 | 0.025626 | 1 | 3.33 | 13.91 | - | - |
| **epithelial cell of proximal tubule** | 685 | 6 | 0.140818 | 1 | 31.29 | 31.53 | - | - |
| **erythrocyte** | 1863 | 6 | 0.924782 | 1 | 12.23 | 35.15 | - | - |
| **fibroblast** | 1386 | 11 | 0.099021 | 1 | 4.05 | 4.33 | - | - |
| **GAT** | 1830 | 6 | 0.916819 | 1 | 7.47 | 18.83 | - | - |
| **granulocyte monocyte progenitor cell** | 5177 | 12 | 0.999877 | 1 | 4.15 | 4.09 | - | - |
| **hematopoietic precursor cell** | 2838 | 10 | 0.930388 | 1 | 6.21 | 4.04 | - | - |
| **hepatocyte** | 2642 | 8 | 0.969086 | 1 | 8.78 | 46.92 | - | - |
| **keratinocyte** | 4020 | 16 | 0.899879 | 1 | 3.74 | 7.51 | - | - |
| **large intestine goblet cell** | 3246 | 14 | 0.798994 | 1 | 3.16 | 6.23 | - | - |
| **late pro-B cell** | 3036 | 8 | 0.991707 | 1 | 2.61 | 5.62 | - | - |
| **luminal epithelial cell of mammary gland** | 1677 | 8 | 0.637735 | 1 | 4.87 | 6.3 | - | - |
| **megakaryocyte-erythroid progenitor cell** | 4553 | 19 | 0.875429 | 1 | 7.25 | 4.76 | 0.0232 | 0.3248 |
| pancreatic ductal cell | 2732 | 19 | 0.102672 | 1 | 13.35 | 7.61 | 0.0496 | 0.6944 |
| pre-natural killer cell | 4359 | 10 | 0.999592 | 1 | 4.29 | 5.73 | - | - |
| **professional antigen presenting cell** | 2856 | 8 | 0.984582 | 1 | 14.58 | 16 | - | - |
| **SCAT** | 1426 | 9 | 0.310103 | 1 | 4.87 | 6.25 | - | - |
| **Slamf1-negative multipotent progenitor cell** | 3527 | 18 | 0.551573 | 1 | 6.2 | 3.75 | 0.0228 | 0.3192 |
| *Slamf1*-positive multipotent progenitor cell | 5421 | 22 | 0.922091 | 1 | 5.35 | 4.25 | 0.1188 | 1 |

| | | | | | | | | |
|---|---|---|---|---|---|---|---|---|
| stem cell of epidermis | 4143 | 14 | 0.975297 | 1 | 3.16 | 4.82 | - | - |
| **B cell** | 529 | 1 | - | - | 4.26 | 5 | - | - |
| **BAT** | 293 | 2 | - | - | 19.14 | 16.49 | - | - |
| **blood cell** | 1387 | 0 | - | - | 0 | 5.44 | - | - |
| **classical monocyte** | 941 | 2 | - | - | 2.61 | 5.49 | - | - |
| **DN1 thymic pro-T cell** | 616 | 1 | - | - | 2.72 | 10.9 | - | - |
| **enterocyte of epithelium of large intestine** | 4178 | 5 | - | - | 5.25 | 9.79 | - | - |
| **granulocyte** | 931 | 2 | - | - | 4.65 | 9.22 | - | - |
| **immature B cell** | 949 | 2 | - | - | 2.49 | 4.24 | - | - |
| **immature natural killer cell** | 664 | 1 | - | - | 4.57 | 13.47 | - | - |
| **immature NK T cell** | 1027 | 3 | - | - | 3.85 | 8.85 | - | - |
| **immature T cell** | 673 | 4 | - | - | 3.56 | 7.57 | - | - |
| **keratinocyte stem cell** | 1328 | 3 | - | - | 4.68 | 6.8 | - | - |
| **kidney collecting duct epithelial cell** | 343 | 0 | - | - | 0 | 42.56 | - | - |
| **Kupffer cell** | 645 | 1 | - | - | 2.49 | 13.02 | - | - |
| **leukocyte** | 987 | 1 | - | - | 3.26 | 5.02 | - | - |
| **lung endothelial cell** | 827 | 3 | - | - | 4.04 | 5.42 | - | - |
| **lung neuroendocrine cells and unknown cells** | 210 | 1 | - | - | 2.43 | 26.29 | - | - |
| **lymphocyte** | 421 | 1 | - | - | 3.36 | 6.19 | - | - |
| **macrophage** | 926 | 0 | - | - | 0 | 6.09 | - | - |
| **MAT** | 427 | 3 | - | - | 7.07 | 13.53 | - | - |
| **mature natural killer cell** | 1161 | 2 | - | - | 3.53 | 10.9 | - | - |
| **microglial cell** | 1021 | 5 | - | - | 3.4 | 6.74 | - | - |
| **monocyte** | 1135 | 1 | - | - | 2.47 | 4.58 | - | - |
| **myeloid cell** | 1240 | 0 | - | - | 0 | 5.85 | - | - |
| **naive B cell** | 583 | 1 | - | - | 4.09 | 4.49 | - | - |
| **natural killer cell** | 643 | 2 | - | - | 2.63 | 11.76 | - | - |
| **pancreatic acinar cell** | 695 | 4 | - | - | 2.97 | 70.68 | - | - |
| **pro-B cell** | 629 | 3 | - | - | 3.18 | 5.99 | - | - |
| **regulatory T cell** | 569 | 1 | - | - | 5.71 | 9.28 | - | - |
| **T cell** | 630 | 4 | - | - | 2.83 | 6.06 | - | - |

### Supplemental Table S4

**Supplemental Table S4.** Imprinted gene enrichment in tissues derived from adult, foetal, embryonic and stem-cell derived cells in the MCA (Han et al., 2018). *Identity* – Tissue identities for the cells used in analysis; All other column descriptions can be found in the legend of Table S2.

| Cell Identity | Up Reg (22,810) | IG (107) | ORA *p* | ORA *q* | Mean FC IG | Mean FC Rest | GSEA *p* | GSEA q |
|---|---|---|---|---|---|---|---|---|
| **Neonatal Muscle** | 1078 | 28 | 6.79E-14 | **2.17E-12** | 4.62244 | 4.211998 | 0.0084 | **0.0168** |
| **Neonatal Rib** | 1223 | 26 | 5.60E-11 | **1.79E-09** | 4.328014 | 4.710165 | - | - |
| **Fetal Stomach** | 470 | 15 | 5.46E-09 | **1.75E-07** | 5.773966 | 5.327665 | 0.2246 | 0.4492 |
| **Neonatal Calvaria** | 1415 | 24 | 2.97E-08 | **9.50E-07** | 5.067572 | 5.561524 | - | - |
| **Placenta** | 1094 | 19 | 7.28E-07 | **2.33E-05** | 9.357343 | 92.72803 | - | - |
| **Embryonic Mesenchyme** | 1013 | 18 | 1.07E-06 | **3.43E-05** | 3.658742 | 4.721681 | - | - |
| **Neonatal Heart** | 1035 | 18 | 1.46E-06 | **4.66E-05** | 3.23287 | 9.304627 | - | - |
| **Pancreas** | 3466 | 36 | 1.58E-06 | **5E-05** | 3.976251 | 6.470602 | - | - |
| **Fetal Intestine** | 622 | 13 | 7.04E-06 | **0.00023** | 3.545314 | 4.465128 | - | - |
| **Neonatal Skin** | 1199 | 17 | 4.18E-05 | **0.00134** | 3.454591 | 5.039246 | - | - |
| **Fetal Kidney** | 792 | 13 | 8.66E-05 | **0.00277** | 2.976385 | 4.847849 | - | - |
| **Fetal Lung** | 484 | 8 | 0.002022 | 0.0647 | 2.889536 | 5.059142 | - | - |
| **Embryonic Stem Cells** | 1277 | 12 | 0.016541 | 0.52932 | 3.438977 | 10.32735 | - | - |
| **Bladder** | 4097 | 27 | 0.03709 | 1 | 3.801455 | 5.669269 | - | - |
| **Male fetal Gonad** | 1328 | 11 | 0.047101 | 1 | 3.119658 | 8.411156 | - | - |
| **Brain** | 3750 | 24 | 0.065314 | 1 | 6.073363 | 13.64887 | - | - |
| **Trophoblast Stem Cells** | 900 | 7 | 0.130384 | 1 | 8.326219 | 5.33632 | - | - |
| **Mammary Gland Involution** | 1104 | 8 | 0.147102 | 1 | 4.604273 | 3.91129 | - | - |
| **Neonatal Brain** | 1506 | 10 | 0.168679 | 1 | 3.343971 | 5.885213 | - | - |
| **Fetal Brain** | 1488 | 9 | 0.26338 | 1 | 4.028713 | 7.870013 | - | - |
| **Kidney** | 1948 | 10 | 0.431517 | 1 | 4.237703 | 39.44992 | - | - |
| **Liver** | 2207 | 11 | 0.463282 | 1 | 3.123224 | 27.4354 | - | - |
| **Uterus** | 3331 | 16 | 0.500819 | 1 | 3.095936 | 5.817416 | - | - |
| **Mouse3T3** | 5910 | 28 | 0.512644 | 1 | 3.686877 | 206.1218 | - | - |
| **Lung** | 1962 | 9 | 0.578215 | 1 | 4.132685 | 11.92437 | - | - |
| **Female fetal Gonad** | 1344 | 6 | 0.608418 | 1 | 13.07576 | 7.853493 | - | - |
| **Bone Marrow cKit** | 2302 | 8 | 0.857716 | 1 | 4.134679 | 3.567175 | - | - |
| **Thymus** | 2110 | 7 | 0.876049 | 1 | 3.314277 | 5.917197 | - | - |
| **Stomach** | 2192 | 7 | 0.899067 | 1 | 4.073947 | 16.60452 | - | - |

| | | | | | | | | |
|---|---|---|---|---|---|---|---|---|
| **Ovary** | 2374 | 7 | 0.937663 | 1 | 3.478591 | 6.889027 | - | - |
| **Small Intestine** | 2227 | 6 | 0.956367 | 1 | 4.713021 | 39.87524 | - | - |
| **Testis** | 4599 | 14 | 0.979018 | 1 | 11.17772 | 8074.858 | - | - |
| **Mammary Gland Pregnancy** | 1418 | 5 | - | - | 3.193922 | 4.501792 | - | - |
| **Muscle** | 1340 | 5 | - | - | 11.96285 | 4.833331 | - | - |
| **Mesenchymal Stem Cells Primary** | 1233 | 5 | - | - | 4.370667 | 8.276373 | - | - |
| **Mammary Gland Virgin** | 1612 | 4 | - | - | 2.704367 | 3.856941 | - | - |
| **Peripheral Blood** | 1552 | 3 | - | - | 3.517534 | 3.867781 | - | - |
| **Fetal Liver** | 1372 | 3 | - | - | 3.543816 | 3.291251 | - | - |
| **Bone Marrow** | 1337 | 3 | - | - | 3.673575 | 3.628222 | - | - |
| **Mesenchymal Stem Cells** | 932 | 3 | - | - | 3.824088 | 4.135905 | - | - |
| **Spleen** | 1989 | 2 | - | - | 3.928225 | 5.671505 | - | - |
| **Prostate** | 442 | 1 | - | - | 3.359508 | 318.2576 | - | - |
| **Mammary Gland Lactation** | 211 | 0 | - | - | 0 | 43.74859 | - | - |

### Supplemental Table S5

**Supplemental Table S5.** Imprinted gene enrichment in global cell types across adult, foetal, embryonic and stem-cell derived tissues in the MCA (Han et al., 2018). *Identity* – Cell identities for the cells used in analysis; All other column descriptions can be found in the legend of Table S2.

| Cell Identity | Up Reg (22,810) | IG (107) | ORA p | ORA q | Mean FC IG | Mean FC Rest | GSEA p |
|---|---|---|---|---|---|---|---|
| **Myocyte** | 807 | 26 | 4.33E-15 | **2.08E-13** | 3.99 | 4.31 | - |
| **Cartilage cell** | 475 | 16 | 7.42E-10 | **3.56E-08** | 7.07 | 8.17 | - |
| **Stromal cell** | 2200 | 31 | 1.39E-08 | **6.67E-07** | 4.08 | 4.62 | - |
| **Chondrocyte** | 1414 | 24 | 2.93E-08 | **1.41E-06** | 5.38 | 7.93 | - |
| **Myoblast** | 854 | 17 | 4.69E-07 | **2.25E-05** | 3.34 | 4.76 | - |
| **Pancreatic acinar cell** | 496 | 12 | 3.74E-06 | **0.0002** | 3.79 | 10.66 | - |
| **Trophoblast progenitor cell** | 1512 | 21 | 6.22E-06 | **0.0003** | 14.73 | 11 | 0.003 |
| **Cardiac muscle cell** | 854 | 14 | 4.60E-05 | **0.0022** | 3.29 | 14.51 | - |
| **Cycling cell** | 458 | 10 | 5.98E-05 | **0.0029** | 3.17 | 4.82 | - |
| **Endothelial cell** | 1549 | 19 | 9.93E-05 | **0.0048** | 3.18 | 5.94 | - |
| **Muscle cell** | 999 | 14 | 0.0002 | **0.0115** | 2.87 | 4.14 | - |
| **Astrocyte** | 2062 | 20 | 0.0013 | 0.0648 | 5.05 | 12.52 | - |
| **Proliferating Myocyte** | 628 | 9 | 0.0028 | 0.1349 | 3.11 | 4.26 | - |
| **Granulosa cell** | 897 | 11 | 0.0033 | 0.1567 | 3.31 | 6.53 | - |
| **Oligodendrocyte** | 3557 | 27 | 0.0065 | 0.3135 | 7.58 | 14.43 | - |
| **Smooth muscle cell** | 1148 | 12 | 0.0076 | 0.3629 | 3.43 | 7.39 | - |
| **Decidual stromal cell** | 1030 | 11 | 0.009 | 0.4316 | 10.53 | 17.66 | - |
| **Endocrine cell** | 1658 | 15 | 0.0105 | 0.5041 | 5.89 | 11.25 | - |
| **mESC** | 1264 | 12 | 0.0154 | 0.7382 | 3.44 | 10.38 | - |
| **Hepatocyte** | 1032 | 10 | 0.0231 | 1 | 4.7 | 33.84 | - |
| **Placenta endodermal cell** | 900 | 9 | 0.0257 | 1 | 4.69 | 12.27 | - |
| **Alveolar type II cell** | 1915 | 15 | 0.0338 | 1 | 3.82 | 8 | - |
| **Proliferative neuron** | 1146 | 10 | 0.0428 | 1 | 3.62 | 5.96 | - |
| **Trophoblast stem cell** | 867 | 8 | 0.0513 | 1 | 7.76 | 5.17 | - |
| **Microglia** | 1115 | 9 | 0.0785 | 1 | 3.24 | 5.52 | - |
| **Neuron** | 1661 | 12 | 0.0889 | 1 | 7.2 | 9.96 | - |
| **Endothelial Cell** | 1741 | 12 | 0.1151 | 1 | 6.47 | 5.49 | - |
| **Spongiotrophoblast** | 896 | 7 | 0.1282 | 1 | 17.55 | 32.32 | - |
| **Schwann cell** | 1268 | 9 | 0.1409 | 1 | 5.62 | 12.47 | - |
| **Kidney epithelial** | 922 | 7 | 0.1425 | 1 | 2.93 | 4.75 | - |
| **Mast cell** | 771 | 6 | 0.1544 | 1 | 7.93 | 9.89 | - |
| **MEF (Cultured)** | 1173 | 8 | 0.1854 | 1 | 3.27 | 4.9 | - |
| **Epithelial cell** | 1203 | 8 | 0.2033 | 1 | 4.17 | 5.58 | - |
| **Proximal tubule brush border cell** | 1984 | 12 | 0.2191 | 1 | 26.02 | 22.44 | - |
| **Cumulus cell** | 1675 | 10 | 0.26 | 1 | 6.64 | 6.28 | - |
| **Osteoblast** | 1068 | 6 | 0.3858 | 1 | 3.82 | 5.8 | - |
| **Urothelium** | 5004 | 25 | 0.3972 | 1 | 4.41 | 8.03 | - |
| **Adipocyte** | 1369 | 7 | 0.4633 | 1 | 4.02 | 6.9 | - |
| **Renal tubular cell** | 1195 | 6 | 0.4921 | 1 | 4.72 | 9.12 | - |
| **3T3 cell** | 5897 | 28 | 0.5073 | 1 | 3.69 | 337.02 | - |
| **Clara Cell** | 1896 | 9 | 0.5358 | 1 | 8.66 | 15.55 | - |
| **Premeiotic germ cell** | 2237 | 10 | 0.6117 | 1 | 9.04 | 5.28 | - |
| **Primordial germ cell** | 1812 | 8 | 0.6229 | 1 | 12.35 | 7.28 | - |
| **Luteal cell** | 3343 | 14 | 0.7179 | 1 | 3.18 | 5.05 | - |
| **Gastric mucosa cell** | 2148 | 8 | 0.8005 | 1 | 3.89 | 6.24 | - |
| **Principal cells of cortical collecting duct** | 3491 | 12 | 0.9099 | 1 | 3.09 | 7.3 | - |
| **Intestinal epithelial cell** | 2411 | 7 | 0.9437 | 1 | 7.8 | 16.98 | - |

| | | | | | | | |
|---|---|---|---|---|---|---|---|
| Testicular cell | 4566 | 14 | 0.9772 | 1 | 11.18 | 8631.25 | - |
| B cell | 1328 | 2 | - | - | 2.88 | 8.02 | - |
| Dendritic cell | 1350 | 1 | - | - | 4.22 | 5.57 | - |
| Eosinophil | 1326 | 4 | - | - | 3.88 | 7.09 | - |
| Erythroblast | 1265 | 1 | - | - | 5.77 | 4.38 | - |
| Erythroid progenitor | 2710 | 3 | - | - | 3.8 | 3.41 | - |
| Intercalated cells of collecting duct | 1055 | 5 | - | - | 2.71 | 11.56 | - |
| Leydig cell | 1098 | 5 | - | - | 3.95 | 5.73 | - |
| Macrophage | 1577 | 2 | - | - | 2.73 | 5.24 | - |
| Monocyte | 1606 | 5 | - | - | 3.03 | 3.97 | - |
| MSC (Cultured) | 915 | 3 | - | - | 3.8 | 4.15 | - |
| Myeloid progenitor cell | 791 | 5 | - | - | 7.58 | 3.65 | - |
| Neutrophil | 1371 | 4 | - | - | 3.62 | 5.47 | - |
| NK cell | 1166 | 2 | - | - | 2.75 | 7.64 | - |
| NK/T cell | 1468 | 5 | - | - | 3.61 | 6.96 | - |
| Prostate gland cell | 439 | 1 | - | - | 3.39 | 290.78 | - |
| Secretory alveoli cell | 217 | 0 | - | - | 0 | 22.44 | - |
| Sertoli cells | 1102 | 5 | - | - | 4.23 | 8.25 | - |
| Stomach epithelial cell | 2033 | 5 | - | - | 4.3 | 9.49 | - |
| T cell | 1716 | 2 | - | - | 3.11 | 6 | - |

### Supplemental Table S6 – Pancreatic Cell Enrichment

**Supplemental Table S6A. Imprinted gene enrichment analysis in pancreatic cells in the MCA (Han et al., 2018).** All column descriptions can be found in the legend of Appendix Table A3.9A

| Pancreatic Cell Identity | Up Reg (9,982) | IG (64) | ORA p | ORA q | Mean FC IG | Mean FC Rest | GSEA p |
|---|---|---|---|---|---|---|---|
| Stromal cell_Smoc2 high(Pancreas) | 977 | 20 | 1.87E-06 | **1.68E-05** | 4.24 | 3.32 | 0.2468 |
| Stromal cell_Fn1 high(Pancreas) | 714 | 13 | 0.001 | **0.005** | 3.44 | 3.54 | - |
| Stromal cell_Mfap4 high(Pancreas) | 600 | 8 | 0.039 | 0.350 | 2.30 | 3.89 | - |
| Endocrine cell(Pancreas) | 393 | 6 | 0.041 | 0.370 | 4.94 | 6.39 | - |
| Î²-cell(Pancreas) | 1088 | 11 | 0.089 | 0.798 | 21.79 | 5.58 | - |
| Endothelial cell_Tm4sf1 high(Pancreas) | 431 | 4 | 0.305 | 1 | 2.23 | 4.40 | - |
| Dividing cell(Pancreas) | 698 | 5 | 0.476 | 1 | 4.88 | 7.81 | - |
| Ductal cell(Pancreas) | 1779 | 12 | 0.489 | 1 | 9.46 | 4.81 | - |
| Endothelial cell_Fabp4 high(Pancreas) | 790 | 4 | 0.764 | 1 | 1.68 | 4.24 | - |
| Acinar cell(Pancreas) | 199 | 2 | - | - | 9.26 | 11.59 | - |
| B cell(Pancreas) | 278 | 2 | - | - | 8.50 | 13.99 | - |
| Dendritic cell(Pancreas) | 384 | 0 | - | - | 0.00 | 6.41 | - |
| Endothelial cell_Lrg1 high(Pancreas) | 362 | 0 | - | - | 0.00 | 6.64 | - |
| Erythroblast_Hbb-bt high(Pancreas) | 373 | 0 | - | - | 0.00 | 9.06 | - |
| Erythroblast_Igkc high(Pancreas) | 158 | 3 | - | - | 5.42 | 7.49 | - |
| Granulocyte(Pancreas) | 401 | 3 | - | - | 16.15 | 16.10 | - |
| Macrophage(Pancreas) | 632 | 1 | - | - | 8.05 | 7.88 | - |
| Macrophage_Ly6c2 high(Pancreas) | 337 | 0 | - | - | 0.00 | 9.21 | - |
| Smooth muscle cell_Acta2 high(Pancreas) | 261 | 2 | - | - | 9.12 | 7.35 | - |
| Smooth muscle cell_Rgs5 high(Pancreas) | 257 | 2 | - | - | 5.11 | 9.01 | - |
| T cell(Pancreas) | 277 | 1 | - | - | 5.85 | 10.27 | - |

**Supplemental Table S6B. Imprinted gene enrichment analysis in pancreatic cells (Baron et al., 2016).** All column descriptions can be found in the legend of Appendix Table A3.9A

| Pancreatic Cell Identity | Up Reg (11,418) | IG (77) | ORA *p* | ORA *q* | Mean FC IG | Mean FC Rest | GSEA *p* | GSEA q |
|---|---|---|---|---|---|---|---|---|
| pancreatic D cell | 4234 | 40 | 0.000593 | **0.004745** | 3.35 | 2.69 | 0.3523 | 1 |
| pancreatic stellate cell | 2067 | 22 | 0.004166 | **0.03333** | 13.72 | 18.68 | - | - |
| endothelial cell | 1883 | 16 | 0.089188 | 0.713501 | 3.79 | 18.29 | - | - |
| pancreatic A cell | 4903 | 34 | 0.164903 | 1 | 2.96 | 1.86 | 0.0619 | 0.1857 |
| pancreatic PP cell | 3224 | 23 | 0.194995 | 1 | 1.66 | 1.86 | - | - |
| pancreatic ductal cell | 3510 | 24 | 0.253517 | 1 | 10.31 | 8.91 | 0.0322 | 0.0966 |
| type B pancreatic cell | 7525 | 48 | 0.284249 | 1 | 2.41 | 2.91 | - | - |
| leukocyte | 1897 | 6 | 0.976333 | 1 | 5.77 | 27.37 | - | - |
| pancreatic acinar cell | 411 | 2 | - | - | 2.38 | 25.78 | - | - |

#### Supplemental Table S7-S9 – Mammary Gland Cell Enrichment

**Supplemental Table S7. Imprinted gene enrichment analysis in the mammary gland at different timepoints in the MCA (Han et al., 2018).** All column descriptions can be found in the legend of Appendix Table A3.9A

| Mammary Gland Timepoint | Up Reg (12,252) | IG (62) | ORA *p* | ORA *q* | Mean FC IG | Mean FC Rest | GSEA *p* | GSEA q |
|---|---|---|---|---|---|---|---|---|
| **Involution** | 3842 | 27 | 0.0287 | 0.0860 | 2.72 | 2.70 | 0.2286 | 0.4572 |
| Lactation | 305 | 1 | - | - | 2.26 | 30.96 | - | - |
| **Pregnancy** | 6213 | 26 | 0.9351 | 1 | 2.89 | 2.31 | 0.1838 | 0.3676 |
| **Virgin** | 4993 | 14 | 0.9992 | 1 | 1.90 | 2.22 | - | - |

**Supplemental Table S8. Imprinted gene enrichment analysis in mammary gland cells different timepoints in the MCA (Han et al., 2018).** All column descriptions can be found in the legend of Appendix Table A3.9A

| Mammary Gland Cell Identity | Up Reg (12,252) | IG (62) | ORA *p* | ORA *q* | Mean FC IG | Mean FC Rest | GSEA *p* | GSEA q |
|---|---|---|---|---|---|---|---|---|
| **Stromal cell(Pregnancy)** | 3260 | 36 | 1.66E-07 | **4.64E-06** | 8.96 | 5.87 | 1.00E-04 | **6.00E-04** |
| **Stromal cell_Col3a1 high(Virgin)** | 1846 | 25 | 1.22E-06 | **3.41E-05** | 7.36 | 4.58 | 0.0099 | 0.0594 |
| **Stromal cell_Pi16 high(Virgin)** | 1282 | 17 | 0.000148 | **0.00413** | 7.04 | 6.92 | 0.3485 | 1 |
| Stromal cell(Involution) | 1243 | 16 | 0.000343 | 0.009599 | 6.18 | 4.95 | 0.1128 | 0.6768 |
| Stromal cell(Lactation) | 133 | 5 | 0.000551 | 0.015427 | 4.14 | 4.25 | - | - |
| **Muscle cell_Pi16 high(Involution)** | 1252 | 15 | 0.001161 | 0.032517 | 9.67 | 6.53 | 0.0882 | 0.5292 |
| Endothelial cell_Fabp4 high(Involution) | 727 | 10 | 0.003302 | 0.092462 | 6.64 | 12.29 | - | - |
| **Muscle cell_Inmt high(Involution)** | 690 | 7 | 0.058614 | 1 | 7.25 | 10.37 | - | - |
| **Luminal cell(Involution)** | 1116 | 9 | 0.107729 | 1 | 5.40 | 6.21 | - | - |
| **Endothelial cell_Aqp1 high(Involution)** | 726 | 6 | 0.159844 | 1 | 16.12 | 14.51 | - | - |
| **B cell_Jchain high(Involution)** | 420 | 4 | 0.162773 | 1 | 7.49 | 12.02 | - | - |
| **Endothelial cell(Pregnancy)** | 636 | 5 | 0.218465 | 1 | 11.62 | 18.07 | - | - |
| **Macrophage_Apoe high(Involution)** | 734 | 5 | 0.313576 | 1 | 7.75 | 7.87 | - | - |
| **Luminal progenitor(Virgin)** | 1170 | 7 | 0.380089 | 1 | 12.51 | 7.64 | - | - |
| **Secretory alveoli cell_Csn2 high(Involution)** | 1225 | 7 | 0.427246 | 1 | 4.38 | 5.93 | - | - |
| **B cell_Cd79a&Fcer2a high(Virgin)** | 685 | 4 | 0.459221 | 1 | 4.45 | 4.25 | - | - |
| **Dendritic cell_Fscn1 high(Pregnancy)** | 710 | 4 | 0.487329 | 1 | 15.08 | 11.50 | - | - |
| **Myoepithelial cell(Pregnancy)** | 2509 | 13 | 0.512325 | 1 | 10.42 | 7.51 | - | - |
| **Dendritic cell_Cst3 high(Virgin)** | 752 | 4 | 0.533244 | 1 | 8.01 | 5.32 | - | - |
| **Macrophage_C1qc high(Virgin)** | 892 | 4 | 0.670297 | 1 | 4.90 | 6.93 | - | - |
| **Secretory alveoli cell_Trf high(Involution)** | 1746 | 8 | 0.674328 | 1 | 5.07 | 4.37 | - | - |
| **Luminal cell_Krt19 high (Virgin)** | 2587 | 12 | 0.681868 | 1 | 6.14 | 5.21 | - | - |
| **Macrophage_Pf4 high(Involution)** | 914 | 4 | 0.689232 | 1 | 5.98 | 6.80 | - | - |
| **Dendritic cell_Fscn1 high(Virgin)** | 1190 | 4 | 0.864731 | 1 | 6.70 | 6.11 | - | - |
| **Macrophage(Pregnancy)** | 2783 | 9 | 0.961082 | 1 | 6.86 | 5.59 | - | - |
| **Secretory alveoli cell(Pregnancy)** | 6973 | 28 | 0.976838 | 1 | 6.01 | 4.49 | 0.0321 | 0.1926 |
| Dividing cell(Virgin) | 2793 | 4 | 0.999878 | 1 | 2.87 | 3.71 | - | - |
| **NK_cells_Gzmb high(Pregnancy)** | 4520 | 9 | 0.999977 | 1 | 3.89 | 4.53 | - | - |

**Supplemental Table S9. Imprinted gene enrichment analysis in the mammary gland at different timepoints (Bach et al., 2017).** All column descriptions can be found in the legend of Appendix Table A3.9A.

| Mammary Gland Timepoint | Up Reg (16,415) | IG (96) | ORA *p* | ORA *q* | Mean FC IG | Mean FC Rest |
|---|---|---|---|---|---|---|
| **Gestation** | 1885 | 16 | 0.0803 | 0.3212 | 5.55 | 7.40 |
| Post Involution | 1062 | 9 | 0.1680 | 0.6719 | 4.14 | 4.81 |
| **Lactation** | 2810 | 20 | 0.1997 | 0.7988 | 5.52 | 11.01 |
| **Nulliparous** | 1464 | 9 | 0.4892 | 1 | 3.06 | 4.22 |
